# Progressive spread of chromosomal inversions blends the role of colonization and evolution in a parallel Galápagos beetle radiation

**DOI:** 10.1101/2022.10.08.511421

**Authors:** Carl Vangestel, Zoë De Corte, Steven M. Van Belleghem, Matthias Vandekerckhove, Karim Gharbi, Frederik Hendrickx

## Abstract

Archipelago island fauna include some of the most compelling examples of parallel adaptation to ecological gradients. However, despite the insuslar nature of these systems, the possibility that repeated instances of within-island divergence resulted from independent (*in situ*) evolutionary rather than repeated colonization events remains most often unclear. Here, we investigated the genomic underpinning of a progressive adaptive radiation of caterpillar-hunter beetles (*Calosoma*) in the low- and highland habitats from the Galápagos. These *Calosoma* beetles have evolved only partially reduced wings in the highland of the youngest islands but evolved to distinct short-winged species in the highland of the oldest islands. In support of independent evolutionary events, the extent of genome-wide divergence between long-winged lowland and short-winged highland populations decreased towards younger islands. However, in support of repeated colonization events, adaptation to highland habitats was driven by repeated selection of alleles that are shared across all highland species and populations. These alleles comprised extensive chromosomal inversions whose origin was traced back to an initial high-lowland divergence event on the oldest island. Moreover, we found evidence that after this initial divergence event, highland alleles spread to younger islands through dispersal of highland individuals as well as dispersal of lowland individuals that were polymorphic at adaptive loci, both providing the opportunity for the establishment of highland populations on the younger islands. These findings highlight the importance of an old divergence in driving repeated adaptation to ecological gradients. Complex histories of colonization and introgression may thus result in a mixed contribution of inter-island dispersal and within-island evolution in shaping parallel species communities on islands.

## Introduction

Insular radiations, like those found on island archipelagos, provide natural laboratories to study the ecological and evolutionary drivers of adaptation and species diversification ^1–3^. Especially when species evolved repeatedly along similar environmental gradients, these cases of parallel within-island evolution have demonstrated that the direction of evolution can be surprisingly predictable ^2^. Moreover, divergence events within islands provide key examples that natural selection can drive rapid speciation in the absence of a geographic barrier ^4,5^ and that niches within an island can either be filled by species that colonized the island as well as through *in situ* radiations ^6–8^. However, as island radiations often include incipient species that still hybridize, the extent to which such repeated *in situ* island radiations represent independent replicates of an evolutionary process or whether they are driven by colonization between islands remains poorly understood ^9–12^.

Independent evolution within islands presumes that the alleles subject to selection evolved independently and, hence, by unique mutations within each island (Fig. 1a) ^13,14^. Yet, an increasing number of studies demonstrate that recurrent ecological differentiation often involves repeated selection on the same alleles ^15–19^, which suggests that a shared evolutionary history and colonization between islands may drive these repeated divergences ^12^. At least three different scenarios, that differ in their degree of gene flow, may explain a shared history of alleles. Although these scenarios can involve a highly different contribution of evolutionary (within-island diversification) versus ecological (colonization) processes in generating ecotypic species pairs, the resulting patterns of genetic differentiation at neutral and adaptive loci can be remarkably similar (Fig. 1) ^20,21^. First, adaptive alleles could be introduced from a nearby island by colonizing individuals that are polymorphic at adaptive loci (“transporter hypothesis”, Fig. 1b) ^22^. Subsequent selection on the alleles at these loci may then result in rapid within-island evolution of similar ecotypes. Second, alleles involved in ecotypic differentiation could be introduced more directly into the gene pool of a resident ecotype through adaptive introgression if few individuals of the alternative ecotype colonize the island and hybridise with the prevalent resident ecotype (Fig. 1c) ^23^. Third, if a larger number of colonizers found a new ecotypic population and hybridize with the resident species, admixture with the resident ecotype may erase the initial genetic differences between these lineages, while maintaining differentiation at loci involved in ecotypic differentiation (Fig. 1d). Under this latter scenario, the repeated occurrence of ecologically similar species on the different islands primarily involves colonization of ecotypes between islands (Fig. 1e) and no within-island diversification is actually involved ^9^. This extent wherein ecotype dispersal between islands have played a role in diversification questions the degree of evidence for parallel evolution for even some of the most iconic examples of adaptive radiation ^9,11^.

**Fig. 1.**
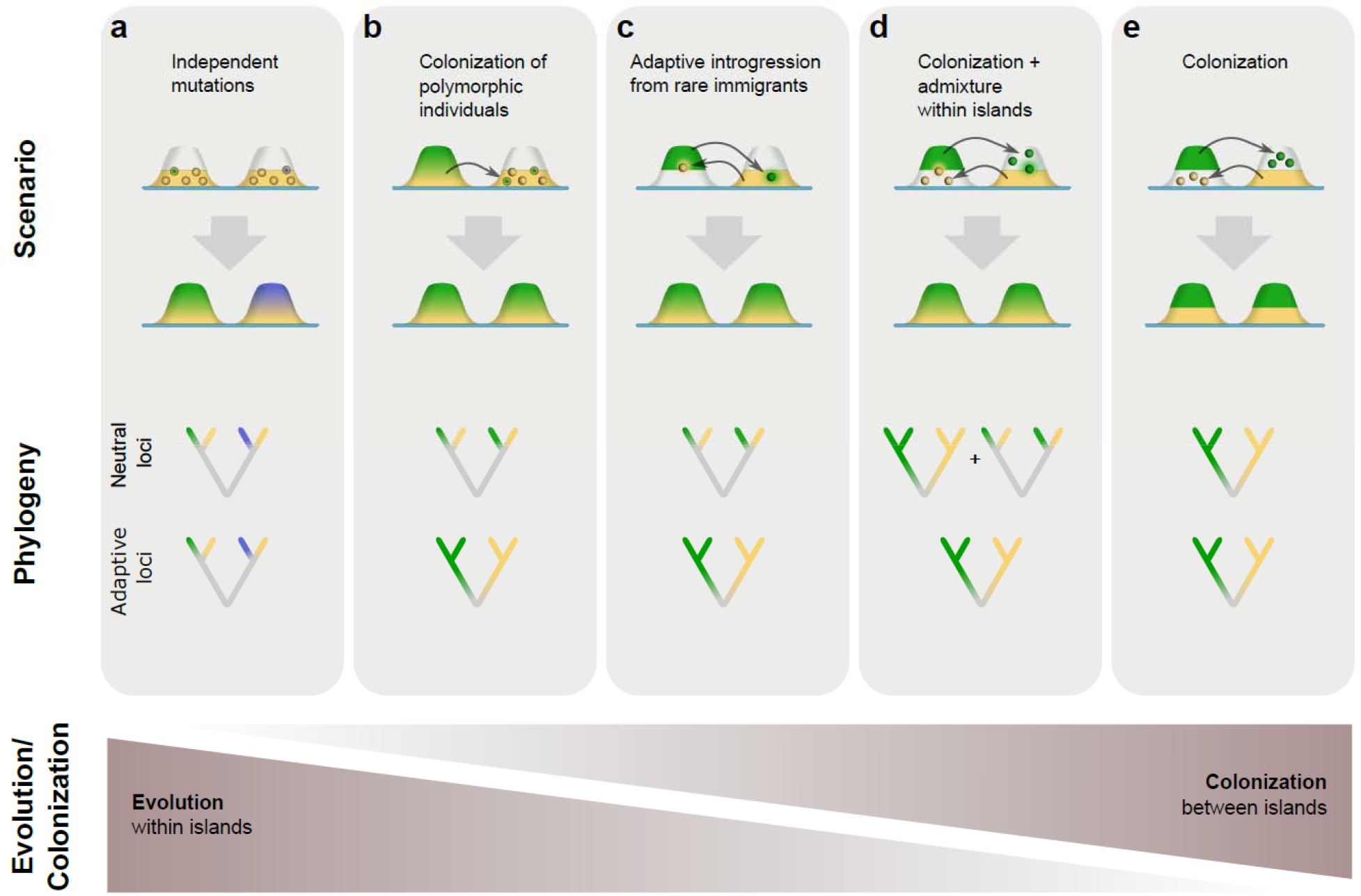
Proposed scenarios resulting in parallel species pairs on islands and involve a gradual contributions of evolution within islands versus colonization between islands. Island areas in yellow are occupied by a hypothetical lowland ecotype, areas in green and blue are occupied by a highland ecotype, white areas represent unoccupied habitat. **a. Repeated evolution by independent mutations**. A unique mutation within each island results in the independent evolution of ecotypes within each island. Ecotypes residing on the same island will be most closely related at both neutral and adaptive loci. **b. Colonization of polymorphic individuals**. Admixture between ecotypes within an island results in polymorphic individuals that colonize other islands. Divergent selection on these polymorphisms results in repeated within-island evolution of both ecotypes. **c. Adaptive introgression from rare immigrants**. Colonization of a single or few individuals of the alternative ecotype that hybridize with the resident ecotype results in adaptive introgression of the alternative alleles. At neutral loci, introgressed haplotypes will be lost by drift due their low frequency. At adaptive loci, selection may maintain introgressed haplotypes and drive the repeated evolution of both ecotypes. **d. Colonization between islands, followed by admixture within islands**. If reproductive isolation between the ecotypes is incomplete, gene-flow within islands may result in a closer relationship between species from the same island at neutral loci. At adaptive loci, introgression is reduced due to divergent selection and maintains the close relationship of ecotypic similar species from different islands. **e. Colonization between islands**. If both ecotypes are reproductively isolated, populations from the same ecotypic species will be most closely related at both neutral as adaptive loci.

Here, we use genome-wide genetic variation to reconstruct the relative contribution of ecological (i.e. colonization between islands) and evolutionary (i.e. *in situ* diversification) processes to a parallel caterpillar hunter beetle (*Calosoma* sp.) radiation in the Galápagos archipelago ^24,25^. The radiation consists of repeated and gradual highland adaptation along the progressive age of the Galápagos islands and we use the different stages of parallel divergence as a powerful solution to infer the role of historical processes and the source of adaptive alleles involved in ecotypic divergence ^26^. At low elevations, a single long-winged species, *C. granatense*, is found on all major islands, while high elevations of the old and intermediate-aged islands San Cristobal (SCB), Santa Cruz (SCZ) and Santiago (SAN) are each occupied by a highland species, taxonomically described as *C. linelli, C. leleuporum and C. galapageum*, respectively (Fig. 2a,b). These highland species share morphological traits such as a marked reduction in wing size, but the degree of morphological divergence with the lowland species decreases towards more recent islands (Fig. 2c) ^25^. Highlands of the youngest islands Isabela (IVA) and Fernandina (FER) are occupied by populations of the lowland species *C. granatense* that evolved a wing size reduction in line with the divergence of the highland species on the older islands (Fig. 2c). We use this system to characterize loci involved in the highland adaptation and leverage the progressive divergence to quantify to what extent ecological versus evolutionary processes underlie parallel adaptation. We do this by first characterizing the phylogenetic incongruencies between loci in this radiation and subsequently studying the evolutionary history of loci that underlie progressive patterns of parallel adaptation. By showing that their parallel divergence was driven by progressive dispersal of highland alleles, encompassing massive chromosomal inversions that evolved during an initial highland divergence on the oldest island, we demonstrate how an initial singular ecotypic divergence and a mixed contribution of colonization and *in-situ* evolution contribute to the emergence of parallel species assemblages in insular systems.

**Fig. 2.**
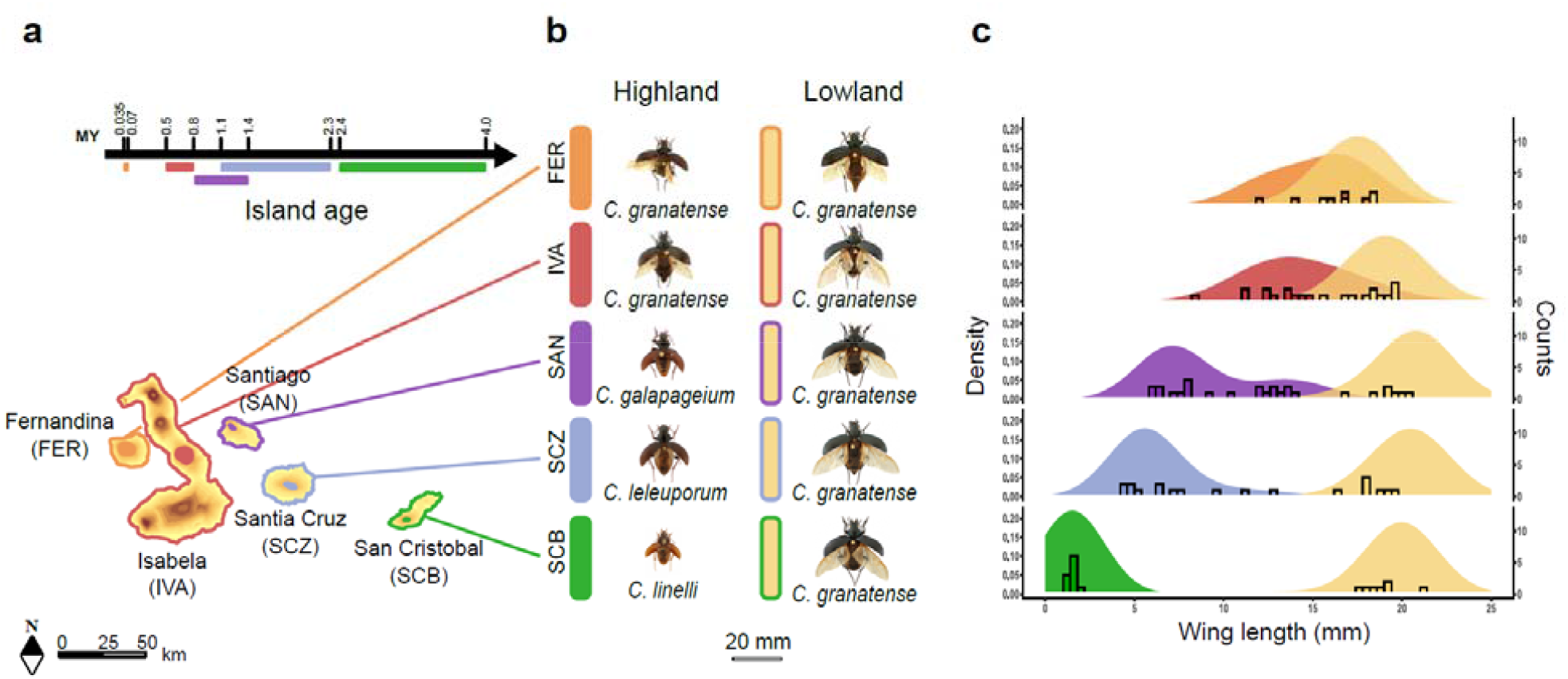
Geographic distribution and wing lengths of Calosoma beetles at the Galápagos islands. **a**. Ages ^32^ of the sampled Galapagos islands. Colored regions show the approximate distribution of the highland species *C. linelli* (green), *C. leleuporum* (light blue), *C. galapageium* (purple), the *C. granatense* highland populations of Volcan Alcedo on Isabella (red) and Fernandina (orange) and the lowland populations of *C. granatense* (yellow). **b**. Pictured specimens of all sampled *Calosoma* species and populations, ordered from the youngest (upper) to oldest (lower) islands. **c**. Density distribution of the wing lengths based on ^24^. Bars show wing lengths of individuals used for genomic analysis in this study.

## Results

### Repeated ecotype divergence strongly correlates with island age

We assessed patterns of genetic differentiation between high- and lowland species from the major islands by mapping 1,135 restriction-site associated DNA sequence tags (RADtags), obtained from 5 to 22 individuals per population, to a newly assembled genome of the lowland species (Table S1, S2 and S3). Average genetic differentiation (*F*_*st*_) between high- and lowland populations was highly consistent with the patterns of morphological divergence and increased almost linearly towards older islands (*r*_*S*_ = 1, *P* = 0.017; Fig. 3a). This increase in genomic high-lowland differentiation was primarily identified as an increased frequency of SNPs with high *F*_*st*_ values. For example, the proportion of highly differentiated SNPs (*F*_*st*_ > 0.4) in the within-island high-lowland comparison increased consistently from 2% on the most recent island Fernandina (FER) to 20% on the oldest island San Cristobal (SCB)(Fig. 3a). A principal coordinate analysis (PCoA) based on SNP genotypes confirmed this gradual differentiation of highland species along the island progression, with the highland species from the oldest island San Cristobal (*C. linelli*, SCB) being the most divergent species, followed by the highland species from Santa Cruz (*C. leleuporum*, SCZ) and subsequently Santiago (*C. galapageium*, SAN) (Fig. 3b).

**Fig. 3.**
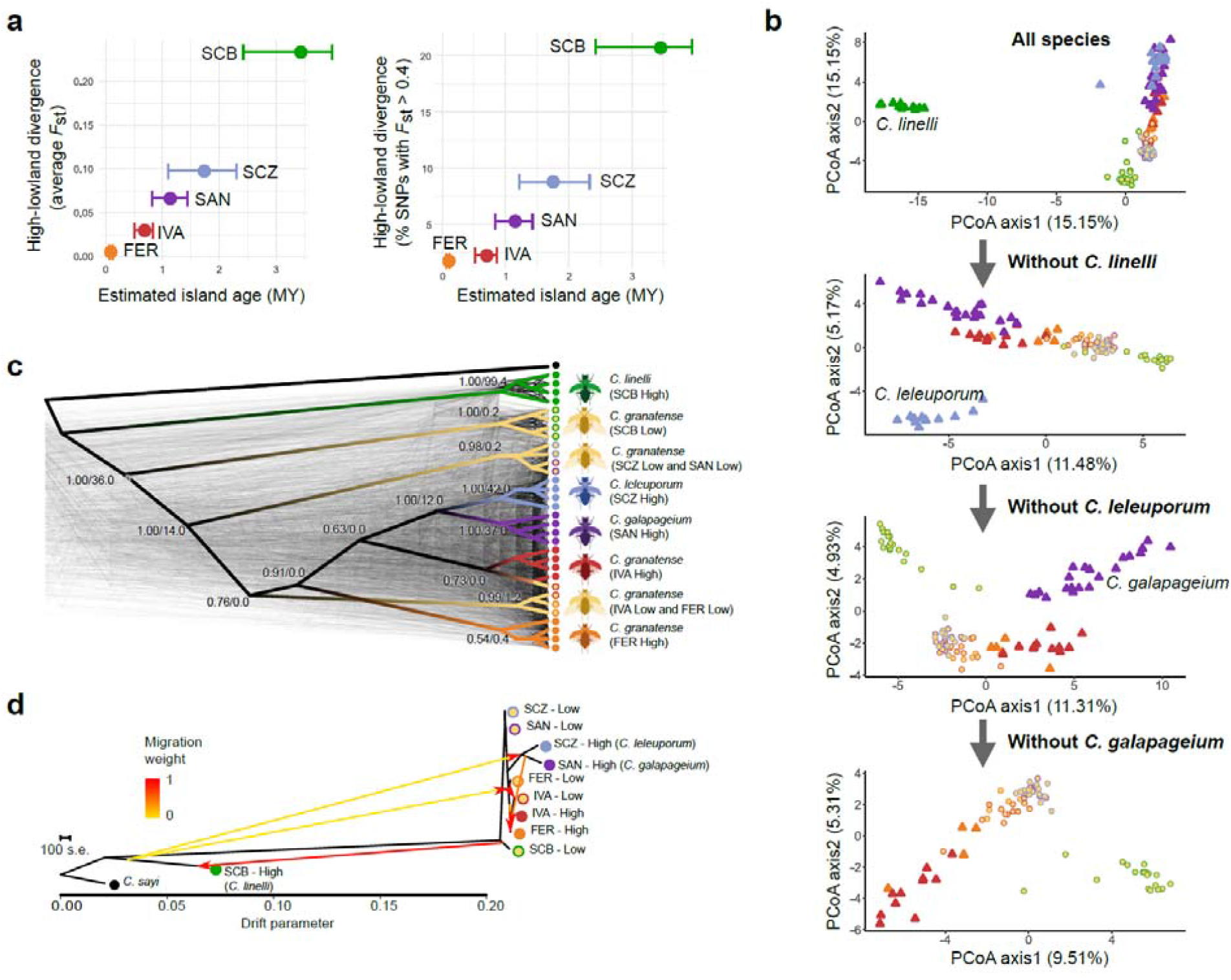
Genetic differentiation and phylogenetic relationships between Calosoma from the Galápagos. **a**. Relationship between island age and within-island high-lowland divergence (left panel: average *F*_st_; right panel: percentage of SNPs with an *F*_st_ > 0.4) and the estimated age of the island. Estimated ages of each island are based on ^32^. **b**. Principal coordinate analysis of the species and populations based on SNP data obtained from RADtag sequencing. The most divergent species was sequentially excluded towards lower panels. **c**. Phylogenetic relationship between the species and populations as inferred from maximum likelihood (ML) trees from 500 random genomic windows of 20kb (light grey trees), with the multispecies phylogeny (ASTRAL ^70^) superimposed. Branch labels show branch support values before slash and gene concordance factors (gCF ^71^) that express the percentage of ML trees containing this branch after the slash. **d**. Population tree and population admixture (arrows) of the different *Calosoma* populations and species, based on SNP data obtained from RADtag sequencing, using TreeMix with six migration edges. Color codes of the different species and populations in all panels are as in Fig. 2.

### Phylogenetic incongruencies suggest admixed species histories

We reconstructed the phylogenetic relationship among the species and populations based on a set of 500 random genomic windows of 20 kb obtained from two (lowland) to four (highland) resequenced individuals per island. Maximum likelihood trees depicted multiple incongruent relationships between the species, illustrating that divergence within this radiation does not follow a clear bifurcating pattern (Fig. 3c). Moreover, a species tree integrating the individual 20 kb window trees supported some basal relationships that were partly incongruent with the patterns of genetic differentiation. The strong divergence between the highland species of the oldest island San Cristobal (*C. linelli*, SCB) from all other species was supported in our phylogenetic analysis as the initial divergence within this radiation. However, the subsequent divergence event split the lowland population of *C. granatense* from this oldest island (San Cristobal) from the remaining species and *C. granatense* populations, which contrasted with the general low levels of genetic differentiation among *C. granatense* lowland populations that we observed in our PCoA analysis based on SNP frequencies (Fig. 3b). The species phylogeny further supported a shared common ancestry of the highland species of Santa Cruz (*C. leleuporum*, SCZ) and Santiago (*C. galapageium*, SAN). The clustering of the highland species from these two islands contrasts strongly with the gradual within-island divergence obtained from the *F*_*st*_ and PCoA analysis where SNP allele frequencies of *C. galapageium* tend to be closer to those of the lowland species *C. granatense* compared with those of *C. leleuporum*. This discrepancy suggests that the highland population of Santiago originated from the highland population of Santa Cruz, but subsequently experienced considerable admixture with the lowland species *C. granatense* after colonization.

Although most nodes in our multilocus species tree were generally strongly supported, gene concordance factors (gCF), which express the percentage of individual trees containing a particular node, were low and ranged from 12% to 36% only, except for the node that clusters the individuals of *C. linelli* (94%) (Fig. 3c). Low gCF values were further noted for nodes that clustered the individuals of the highland species *C. leleuporum* (42%) and *C. galapageium* (37%), indicating that these species share a substantial proportion of their haplotypes with other species for most of the investigated loci.

The tree topology as inferred from TreeMix ^27^ confirmed the strong divergence of the highland species of San Cristobal (*C. linelli*) from the closely related remaining species and populations (Fig. 3d). Congruent with our multispecies phylogeny, the remaining two highland species (*C. leleuporum* and *C. galapageium* of Santa Cruz and Santiago respectively) grouped in a clade that was situated within the populations of the lowland species *C. granatense*. Both TreeMix and an introgression analysis based on f_4_ statistics ^28^ supported numerous interspecific migration events within as well as between islands (Supplementary Methods 1). Ancestral genetic variation of the distinct highland species of San Cristobal (*C. linelli*) was retained in the *C. granatense* lowland population of this island as well as in the two other highland species (*C. leleuporum* and *C. galapageium*). Signatures of such ancient admixture could even be traced back to the highland populations of *C. granatense* inhabiting the youngest islands (Fig. 3d, Supplementary Methods 1, Table S4).

### Outlier loci are shared across islands

Within islands, we identified the genomic regions associated with high-lowland divergence by screening the RADtag sequences for SNPs that were significantly more differentiated compared to background levels in the within-island comparisons (BayeScan ^29^, Q-value < 0.01). We identified between 196 (Isabella, IVA) and 458 (Santiago, SAN) outlier SNPs in these separate within-island comparisons, except for the most recent island Fernandina for which a lower number of individuals could be sampled (Fig. 4b). These outlier SNPs were not randomly distributed across the genome, but generally clustered into large genomic blocks extending up to several megabases (Fig. 4b). Genomic regions characterized by an elevated divergence in the high-lowland comparison within each island were highly consistent across the different islands, indicating that largely the same genomic regions are involved in high-lowland divergence in the different species and islands.

**Fig. 4.**
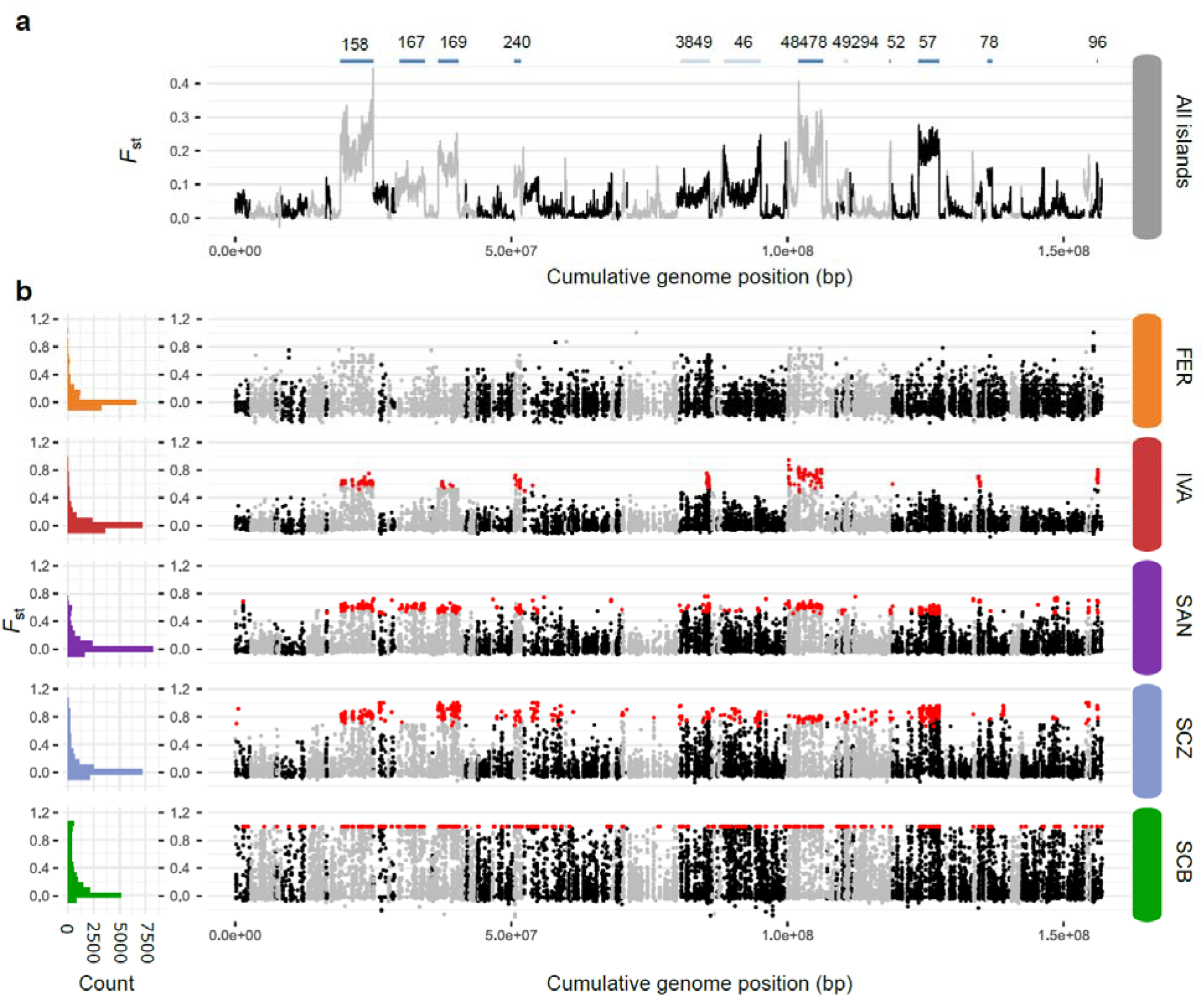
Genetic differentiation between the high- and lowland species or populations. **a**. Genetic differentiation (*F*_st_) between all resequenced high-versus lowland individuals (20kb windows). Upper blue line segments show the location of genomic regions tested for the presence of structural variations (SV), with those in dark blue being regions showing support for SV based on anomalies in the orientation and insert size of read mappings as detected by *BreakDancer* ^72^. Scaffold ID’s are given above each SV. **b**. Differentiation (*F*_st_) between high-and lowland individuals within each island at individual SNPs obtained from RADtag sequencing. Left histograms show the *F*_st_ frequency distribution across the entire genome and right panels shows their location at the different genomic scaffolds. SNPs indicated in red are outlier SNPs with a significantly higher differentiation than expected by chance within each island comparison (BayeScan ^29^; Q-value < 0.1).

### Outlier loci include extensive chromosomal inversions

Because most outlier SNPs were concentrated into large genomic blocks that were shared across islands, we tested if the alleles under divergent selection potentially included structural variations (SVs). We investigated the presence of such SVs for the twelve longest contiguous genomic regions with elevated divergence (*F*_*st*_ > 0.1) in an overall high-lowland comparison (Fig. 4a). Each region contained at least one RADtag with a SNP identified as outlier in a minimum of two within-island ecotype comparisons, with a total of 109 outlier RADtags (57% of all identified outlier RADtags) across all twelve regions combined, which supports their association with high-lowland divergence (Fig. 4, Table S5). An SV analysis based on anomalies in the orientation and insert size of read pairs identified chromosomal inversions that perfectly overlapped with the five largest (3.7 to 5.9 Mb) and one smaller (213 kb) region of elevated divergence (Fig. 5a, Fig. S1, Table S5). Two additional regions of elevated divergence included the scaffold start and read pairs situated at the potential SV breakpoint mapped to another scaffold and thus likely represent partially assembled inversions. One last region was flanked by deletions and potentially comprises a more complex SV. Three remaining regions did not show evidence for SV based on anomalous read mappings at the flanking regions (Table S5).

**Fig. 5.**
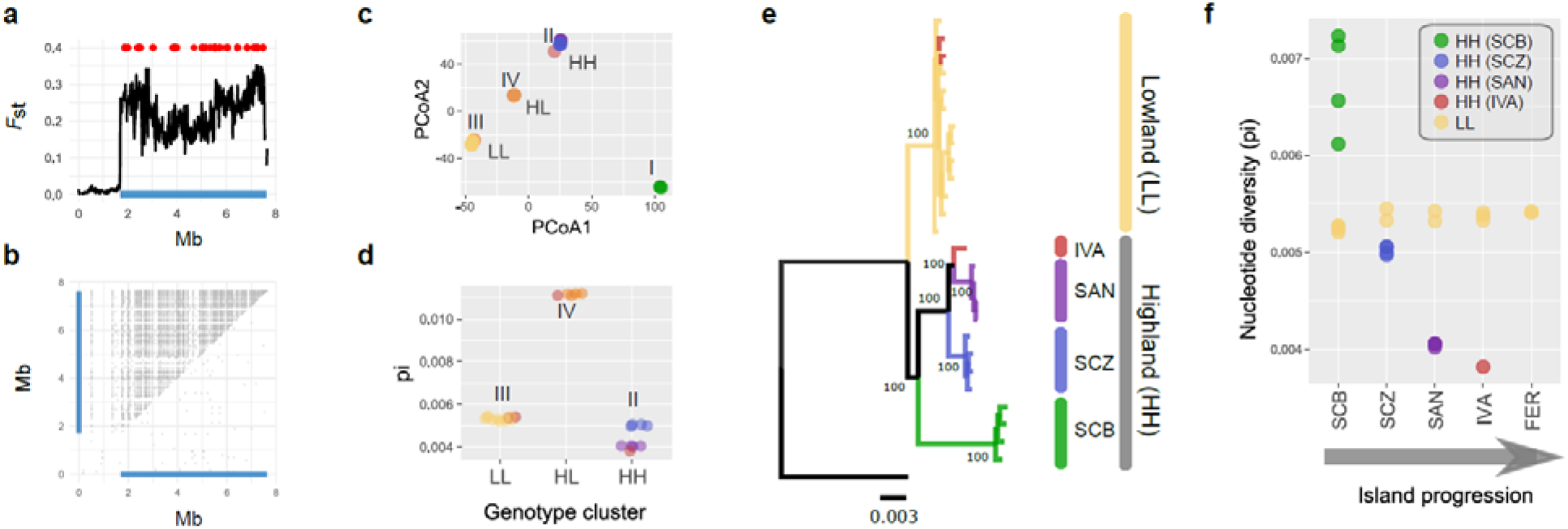
Structural variations (SV) underlie distinct alleles associated with high-lowland divergence. Results of a single scaffold (scaffold158) are shown. Results for the other scaffolds with SV associated with high-lowland divergence are shown in Fig. S1. **a**. *F*_ST_ distribution (20kb windows) based on a comparison between all resequenced high-versus lowland individuals. Blue bottom line shows the location of the chromosomal rearrangements detected by *BreakDancer*^72^. Red dots show the location of RADtags identified as outlier loci in at least one within-island highland-lowland comparison. **b**. Location of SNPs in perfect linkage disequilibrium (*r*^2^ = 1). Grey dots above diagonal show *r*^2^ = 1 values for all 32 resequenced individuals. Grey dots below diagonal show ^*r*2^ = 1 values for homozygous LL individuals only. **c**. PCoA based on SNPs located at the inversion (blue line in panel a). HH, LL and HL refer to the cluster of individuals genotyped at the inversion as homozygous for the highland allele (HH), lowland allele (LL) and heterozygous (HL). **d**. Differences in nucleotide diverssity at the inversion between individuals genotyped as HH, HL and LL in panel c. **e**. Maximum likelihood tree of the nucleotide sequence at the inversion. Individuals genotyped as heterozygotes (HL) were excluded from the analysis. Node values represent bootstrap values based on 1000 replicates. The tree was rooted with the mainland species *C. sayi*. **f**. Relationship between individual nucleotide diversity at the inversion and the progression of the islands. Only individuals genotyped as homozygous for the lowland allele (LL, yellow) and highland allele (HH, remaining colors) are included. Color codes are as in Fig. 2.

SNPs located on each SV generally showed an identical segregation pattern with no obvious decay in linkage disequilibrium across the entire length of the SV (Fig. 5b, Fig. S1 and S2), providing additional support that extensive non-recombining and highly divergent haplotypes underlie these regions of elevated divergence. A local PCoA analysis based on the SNP genotypes within each SV consistently clustered individuals into four distinct groups (Fig. 5c, Fig. S1): (I) a cluster of individuals from the highly divergent species *C. linelli*, (II) a cluster of individuals from the highland species *C. leleuporum, C. galapageium* and the highland populations of *C. granatense*, (III) a cluster that mainly comprised individuals from the lowland species *C. granatense* and (IV) a cluster containing individuals of both high-and lowland species and populations that was situated in-between clusters II and III. This clustering is consistent with the presence of a distinct high (H) - and lowland (L) allele, with clusters II and III comprising individuals homozygous for the high- and lowland allele, respectively, and cluster IV corresponding to individuals being heterozygous for both alleles. Heterozygosity for a distinct high- and lowland allele in the individuals of this latter cluster (IV) was supported by their markedly higher nucleotide diversity at the SV compared with those homozygous for one of the two alleles (clusters II and III, Fig. 5d). Moreover, these elevated levels of nucleotide diversity in heterozygotes were maintained across the entire length of the SV, which provides additional support for the lack of recombination among those divergent haplotypes (Fig. S2).

### Chromosomal inversions originated from an initial high-lowland divergence and progressively spread towards younger islands

To infer the evolutionary history of the chromosomal inversions, we constructed maximum likelihood phylogenies of the haplotypes present at each SV. Haplotypes associated with all highland species and populations, including the highly distinct haplotypes of the most divergent highland species *C. linelli* from San Cristobal, consistently clustered with high support into a monophyletic clade (Fig. 5e, Fig. S1). Thus, haplotypes selected in all highland species and populations appear to have a single evolutionary origin and subsequently spread across all islands. Spread of highland alleles generally followed the island progression as shown by a consecutive split of highland haplotypes according to island age for six SVs (Fig. 5e, Fig. S1). This progressive spread was further corroborated by a significant and consistent decrease in nucleotide diversity of highland alleles towards younger islands for all but one SV (Fig. 5f, Fig. S1, Table S5). The highly consistent phylogenetic and nucleotide diversity patterns across the different SV could, at least partially, be caused by a tight physical linkage of the scaffolds on which the SV are located. However, none of the SV showed an identical segregation pattern across the 32 investigated individuals, which demonstrates that they represent independently evolved loci with a shared evolutionary history (Fig. S3).

Finally, we investigated if the divergence between the high- and lowland associated alleles across all SVs evolved during a singular high-lowland divergence event by comparing their timing of the split between high- and lowland alleles (Fig. 6a). Estimated divergence times, expressed relative to the divergence from the mainland species *C. sayi*, ranged between 0.5 and 0.75 and 95% posterior density intervals of the splitting times overlapped for six of the nine SV. These estimated divergence times were centered around the estimated divergence time of the most ancient high-lowland divergence in our species phylogeny, being the split that gave rise to the highland species *C. linelli* and the remaining species (Fig. 6b). This suggests that the evolution of highland alleles coincides with the most ancient high-lowland divergence in this radiation.

**Fig. 6.**
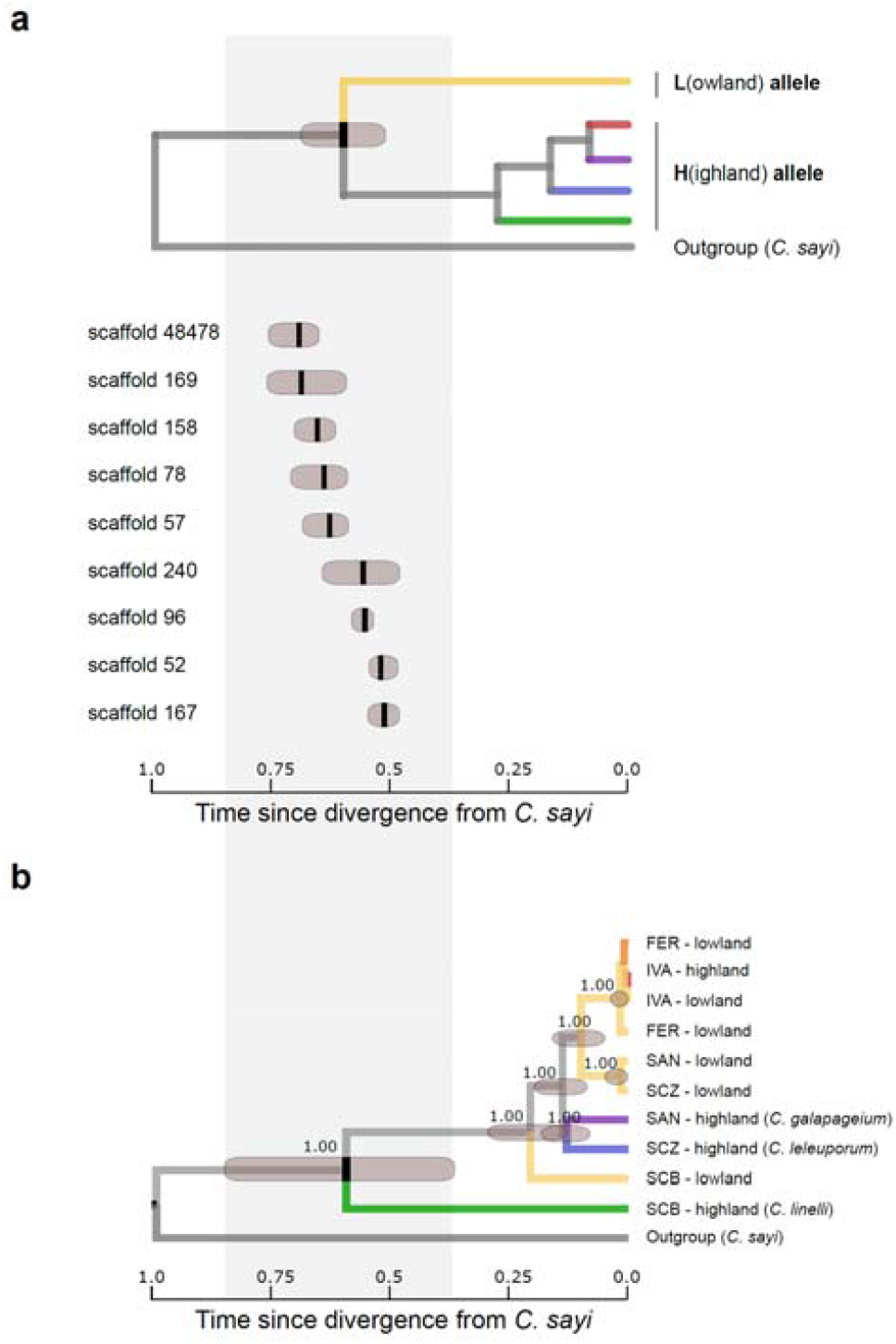
Relative divergence time between high- and lowland alleles for nine different SV compared to the relative divergence times of the species and populations. **a**. Estimated divergence time (black vertical bars: mean; dark grey bar: 95% highest posterior density) between high- and lowland alleles for nine different structural variations (SV) associated with high-lowland divergence. **b**. Estimated divergence times of the different species and populations based on random selection of 50 genomic windows of 20kb that are located outside the SV. Node values represent posterior probabilities of the clades. Divergence times in both panels are scaled to the same time scale, expressed as relative time since the divergence from the mainland species *C. sayi*.

## Discussion

Archipelagos where islands emerged following a known chronosequence have proven key to reconstruct the processes and mechanisms of species diversification ^2,3,30^. Here, we inferred the genomic basis of a gradual divergence of multiple highland species and populations from a single lowland species along the progression of the Galápagos islands. Patterns of genome-wide divergence corroborated the lower high-lowland divergence towards younger islands, which hints towards a gradual, repeated, and independent within-island evolution of distinct highland species and populations on the different islands. However, adaptation towards highland habitats largely involved selection within the same genomic regions across all highland species. These genomic regions were characterized by extensive chromosomal inversions, often extending multiple megabases, that resulted in distinct and non-recombining haplotypes associated with high-lowland divergence. The well-supported monophyletic clustering of highland associated haplotypes provides strong evidence that a single evolutionary origin underlies the highland alleles at each locus. Moreover, high- and lowland selected alleles diverged within a similar time frame across loci that corresponded with the estimated split time of the most anciently diverged highland species *C. linelli* from San Cristobal. Thus, the repeated evolution of highland species and more recent highland populations appears driven by selection on alleles that evolved during a singular high-lowland divergence event that can be traced back to the most ancient high-lowland species divergence within the archipelago.

The phylogenetic pattern of haplotypes that were associated with highland habitats depicted consecutive splits consistent with the chronosequence of the islands. This consistent phylogenetic signal was likely preserved by the association of highland haplotypes with reduced dispersal capacity, which increased their geographic isolation and hampered their subsequent exchange between the islands ^31^. Combined with a consistent decrease in nucleotide variation towards younger islands, consistent with serial and stepwise founder events of highland alleles, strongly supports a progressive spread of highland alleles along this island progression. Propagation of highland alleles towards more recent islands might either have occurred through (i) dispersal of highland species between islands (Fig. 1d), (ii) dispersal of few highland individuals that introduced highland alleles in resident lowland populations (‘adaptive introgression’ ^23^, Fig. 1c) or (iii) dispersal of lowland individuals that are polymorphic at loci involved in high-lowland divergence (Fig. 1b). Our data suggest that several of these mechanisms took place within this radiation. Dispersal of the highland species towards younger islands was supported by the sister relationship of the highland species of Santa Cruz (*C. leleuporum*) and Santiago (*C. galapageium*) in our phylogenetic analysis, which indicates that the highland species of Santiago evolved from highland individuals of Santa Cruz that colonized Santiago, rather than through an *in-situ* high-lowland divergence on Santiago. This appears plausible given the connection between both islands about 1 Mya ^32^. However, subsequent gene-flow with the lowland species *C. granatense* took place after this colonization event and almost completely erased the phylogenetic signature of the sister relationship between those two highland species. Our genome-wide phylogeny revealed that only 12% of the genome (gCF) still supports the sister relationship between these highland species on the youngest islands. In contrast, spread of highland alleles by dispersal of polymorphic individuals of the lowland species *C. granatense* that likely resulted in the introduction of highland alleles took place between, at least, the more recent islands Isabela and, particularly, Fernandina. For these islands, we found that winged lowland individuals of *C. granatense* are polymorphic at loci under divergent selection (Fig. S3), which may easily result in the transport of highland alleles between islands. Dispersal of ‘pure’ highland individuals between these islands can moreover be excluded as highland habitats of these more recent islands are not populated by distinct highland species but by individuals of the lowland species with reduced wings and a higher frequency of highland alleles.

While decreasing levels of ecotypic divergence along a chronosequence is often considered to represent different stages of the speciation process ^26,33^, our results rather point towards a reversal of the high-lowland divergence along this island progression. More precisely, while high- and lowland species are morphologically and genetically well differentiated on the oldest island San Cristobal, their divergence decreases towards the younger islands Santa Cruz and Santiago. Provided the sister relationship of the highland species on these latter two islands, this lower genomic divergence on Santiago implies increased admixture from the lowland species into the highland on this younger island (Fig. 1d). Increased admixture between high- and lowland species on more recent islands may ensue from the stepwise colonization of highland species on islands with a resident lowland population. This leads to decreasing founder population sizes of highland species on more recent islands, which on its turn increases asymmetric introgression from the large population of the resident lowland species *C. granatense* into the smaller colonizing population of highland individuals ^34,35^. If the number of immigrant highland individuals becomes very small, these individuals may even be more likely to exchange genes with the resident lowland population rather than with other highland immigrants, resulting in adaptive introgression ^23^ of highland alleles in the resident lowland population (Fig. 1c). Selection on these introgressed alleles in highland populations may then result in a repeated evolution of the highland ecotype. Because introgression of adaptive alleles leads to polymorphisms at adaptive loci, highland alleles may ultimately be introduced by the immigration of such polymorphic individuals (Fig. 1b), a process that likely took place at the youngest islands Isabela and Fernandina of the archipelago. Therefore, patterns of divergence along this island progression provide unique support and an extant illustration of the presumed stages of the emerging ‘two-time frames’ model of repeated ecotype evolution, which proposes that contemporary ecotypic evolution is driven by selection on alleles that potentially originate from an old, singular and even allopatric ecotypic divergence ^16,17^.

## Conclusion

The repeated occurrence of parallel species pairs on insular systems is presumed to result either from the colonization of species between islands or through repeated *in-situ* adaptation ^2^. Results from our study demonstrate that complex introgression patterns between as well as within islands challenge this dichotomous view and suggest that the difference between these two mechanisms is likely more gradual as generally assumed. Habitat patches on islands are highly dynamic with respect to their spatial and temporal distribution, often driven by climatic and geological dynamics, resulting in multiple episodes of fission and fusion between ecotypes ^30,36^. This may both erase the original phylogenetic signals of species divergence and result in the exchange of alleles involved in ecotypic differentiation between islands. Chromosomal rearrangements could strongly facilitate the repeated evolution after adaptive introgression and resist the effect of homogenizing gene-flow by maintaining favorable allelic combinations ^37–41^. Given the frequently reported evidence of interspecific gene-flow in island radiations like for example Darwin finches ^42,43^, giant tortoises ^36,44^, *Hogna* wolf spiders ^45^ and Hawaiian *Metrisoderos* trees ^18^, these complex introgression patterns can be expected to be ubiquitous. The number of founding individuals or haplotypes of the alternative ecotype that colonize an island and the amount of interspecific gene flow between the two ecotypes after colonization are likely key factors that determine the relative contribution of inter-island colonization and within-island diversification in the origin of parallel species assemblages on islands. Better comprehending the interplay between these mechanisms could help to better predict how rates of colonization and speciation determine biodiversity dynamics on islands ^12,46^.

## Methods

### Sampling

We sampled all high- and lowland species or populations from all islands for which distinct highland ecotypes or species have been reported at the Galapagos archipelago i.e. San Cristobal (SCB), Santa Cruz (SCZ), Santiago (SAN), Isabela (IVA) and Fernandina (FER) (Table S1, Fig 2). The island Isabela is the only island within the Galapagos that comprises multiple large volcanoes and we restricted our sampling to the most centrally located Volcan Alcedo. Individuals were sampled during different sampling campaigns between 1996 and 2014 (Table S1) and stored live in liquid nitrogen or pure ethanol shortly after sampling. While high- and lowland species are easily identified in the field on the oldest islands San Cristobal, Santa Cruz and to a lesser extent Santiago ^24^, divergence of highland ecotypes varies more gradually on the younger islands Isabela and Fernandina. To ensure that individuals of the highland ecotype were sampled at these islands, only those individuals sampled at the outermost volcano summit (1110m and 1290m at Isabela and Fernandina respectively) were considered as highland ecotypes. We included between 11 and 22 individuals per population for genetic analysis, except for Fernandina where only five individuals of the high- and lowland population could be sampled. Wing-sizes of the sampled individuals clearly matched the earlier reported wing sizes of these species and populations (Fig. 2b,c) and confirmed the gradual reduction in wing-size of the highland species and populations towards younger islands ^47^. For the most recent island Fernandina wing sizes of highland individuals overlapped with those from lowland individuals. We further sequenced the genome of a single specimen of the related mainland species *Calosoma sayi* ^48,49^, sampled by M. Husemann in Texas, USA, and was used as an outgroup species. *C. sayi* is one of the most closely related species with those found at the Galapagos and taxonomically classified within the same subgenus *Castrida* ^50^.

### Genome assembly

We assembled the genome of *Calosoma granatense* using both paired-end libraries with short insert sizes of 170bp, 500bp and 800bp and mate-paired libraries with insert sizes of 2kb, 5kb, 10kb and 20kb. Short-insert size libraries were all constructed from a single individual sampled at Santa Cruz at 350m altitude, while long-insert mate paired libraries were constructed from DNA extracts from nine different individuals that all originated from this same locality (Table S1). Total DNA was extracted from these individuals with the NucleoSpin ® Tissue Kit, Macherey-Nagel GmBH and library construction and sequencing performed at the Bejing Genomic Institute, Hongkong. Sequencing errors were corrected based on the *k*-mer frequency spectrum with SOAPec ^51^, specifying a *k*-mer value of 17. Corrected reads were then used as input for genome assembly with Platanus^52^ using default settings. Contigs were constructed based on the short-insert libraries only with the ‘platanus assemble’ tool, and subsequently combined into scaffolds with the ‘platanus scaffold’ tool using both short- and long-insert libraries. Gaps between the scaffolds were finally filled with the ‘platanus gap_close’ tool using both short- and long-insert libraries. The final assembly consisted of 6045 scaffolds summing to a size of 167,880,245bp (Table S3). We estimated the genome size by obtaining the *k*-mer frequency spectrum from whole genome sequencing data from multiple individuals (see ‘*Whole genome resequencing*’ in ‘*Methods*’) with Jellyfish v2.3.0 ^53^ and analyzed the frequency distribution with GenomeScope ^54^. These analyses yielded an estimated genome size of 173.3 Mb (± 6.0 SD) for a *k*-mer = 21 and highly similar estimates for other tested *k*-mer sizes (*k*-mer= 17: 172.6 Mb ± 8.2SD; *k*-mer = 31 : 172.6 Mb ± 8.2SD). Based on these estimates, the assembled genome represents 97% of the estimated genome size. Completeness of the assembly was further assessed based on a set of 1,658 benchmarked single copy orthologs (BUSCO’s) from Insecta ^55^. Screening the draft genome for these BUSCO’s revealed that 85.3% were present in our assembly, of which 0.3% duplicated and 8.4% fragmented (Table S3). We screened the genome for repetitive elements with RepeatMasker v1.295 ^56^ specifying ‘Coleoptera’ as species and constructed a library of *de novo* repetitive elements with RepeatScout v1.0.5^57^. Both methods resulted in a total repeat content of 11.04% (Table S3).

### Restriction-site associated sequencing (RADseq) and whole genome resequencing

We performed RADseq on between 11 and 22 individuals of each population, except for the island of Fernandina where only five individuals of the high- and lowland population were available (Table S1). DNA was extracted using the NucleoSpin ® Tissue Kit, Macherey-Nagel GmBH following the manufacturer’s instructions. DNA extracts were normalized to a concentration of 7.14ng/µL and RADtag libraries were constructed following the protocol described in ^58^ using the SbfI-HF restriction enzyme (NEB) and sequenced on either an Illumina MiSeq (2×250bp) or HiSeq1500 (2×100bp) platform. Raw data were demultiplexed to individual samples using the *process_radtags* module in Stacks v1.20 ^59^. PCR duplicates were removed with the *clone_filter* tool based on identical reverse read ends. Paired reads were mapped to the draft reference genome with BWA (bwa mem) ^60^ using default settings and SNPs were called using GATK’s UnifiedGenotyper tool. Only biallelic SNPs (--max-alleles 2) with a minimal SNP quality (--minQ) of 60, an individual genotype (--minGQ) quality of 30 in at least 80% of the individuals (--max-missing) and a minimum allele frequency (--maf) of > 0.05 were retained with VCFtools ^61^. We further excluded all positions located in repetitive regions detected by RepeatMasker v1.295 ^56^ (see *Genome assembly*). After filtering we retained 15,256 SNPs.

We further sequenced the genomes of 33 individuals comprising four individuals of each highland species and population, four individuals of the *C. granatense* lowland population of San Cristobal, two individuals of the remaining *C. granatense* lowland populations and an individual of the related species *C. sayi* from Texas, USA (Table S1 and S2). Genomic libraries were constructed with the TruSeq Nano DNA LT kit (Illumina) following the manufacturer’s instructions and sequenced on an Illumina HiSeq1500 platform (2×100bp). Resulting sequencing reads were mapped to the draft reference genome with BWA (bwa mem) ^60^ with default settings. Local indel-realignment was performed using GATK’s RealignerTargetCreator and IndelRealigner ^62^. Variants were first called for each individual sample using GATK’s HaplotypeCaller and subsequently called across all samples with the GenotypeGVCF tool. Variants were finally hard filtered with the VariantFiltration tool specifying the following five criteria: quality score normalized allele depth (QD) < 2.0, FisherStrand (FS) > 60.0, MappingQuality (MQ) < 40, Mapping Quality Rank Sum (MQRandSum) < -12.5 and ReadPosRankSum < -8.0 and removed SNPs located in repetitive regions. A total of 15,569,155 SNPs were retained after filtering, of which 98% were shared by at least 80% of the resequenced individuals.

### Patterns of genomic divergence

We estimated genome-wide genetic differentiation between the high- and lowland populations within each island based on the *F*ST-values of SNPs obtained from our RADseq data. The genetic population structure between high- and lowland individuals was further explored with a principal coordinate analysis (PCoA) using the R package adegenet 2.1.3 ^63^. To minimize linkage disequilibrium between SNPs we randomly selected a single SNP within each RADtag (n=1135) using an in-house Python script. We sequentially discarded the most diverged species in subsequent PCoA’s to further reveal the hierarchical population structure and to explore the relationships among the remaining clusters in more detail.

To assess whether these SNPs were a putative target of natural selection we evaluated whether among-population genetic differentiation was significantly higher than expected under neutrality using BayeScan v2.1 ^29^. This outlier analysis calculates the posterior probability of each SNP to be the target of selection by contrasting a model which includes the effect of selection to one excluding such effect. Simulations were performed using a total of 20 pilot runs of 5000 iterations each to tune model parameters. Subsequently we ran the Markov chain Monte Carlo for another 100 000 iterations, discarded the first 50 000 as a burn-in while setting the prior odds for the neutral model to 10 and used the internal q-value function of the software package to assess significance at a FDR threshold of 0.1 (q<0.1).

Genetic differentiation between all resequenced high- and lowland individuals was assessed using a sliding window approach to minimize noise from SNP based divergence estimates. Weir and Cockerham’s *F*ST statistics ^64^ were estimated for non-overlapping 20 kb windows using VCFtools v0.1.16 ^61^.

### Phylogenetic analysis

We inferred the phylogenetic relationship between the resequenced individuals based on 500 windows of 20kb each. Windows were selected with a custom python script (SelectRandomWindows.py) and written to gff format specifying the start and end-positions of each window. Fasta files containing the individual sequences for each window were then extracted with the vcf2fasta.pl tool (https://github.com/santiagosnchez/vcf2fasta), using the genome assembly, genome wide VCF and gff-file specifying the locations of the 20kb windows as input files. We estimated maximum likelihood trees for each window-specific fasta with IQ-TREE ^65^, specifying 1000 ultra-fast bootstrap samplings ^66^. Before tree estimation, best substitution models for each window were selected using ModelFinder ^67^ as implemented in IQ-TREE. ML trees were visualized with the densiTree function implemented in the *phangorn* v2.5.5 ^68^ package in R v4.0.3. To ease visualization, trees were first made ultrametric by transforming the branches proportionally in FigTree v1.4.2 (http://tree.bio.ed.ac.uk/software/figtree). We then reconstructed a multispecies phylogeny that accounts for the potential discordance in the window-specific phylogenies using ASTRAL ^69,70^. Besides the calculation of branch support values as implemented in ASTRAL ^70^, we also calculated gene concordance factors (gCF) ^71^, which express the percentage of gene (window) trees containing this branch. gCF’s were obtained from IQ-TREE by specifying the multi-species consensus tree from ASTRAL as reference tree. We applied a graph-based model implemented in TreeMix v1.13 ^27^ to explore evolutionary relationships and admixture events among resequenced high- and lowland populations and species. A maximum-likelihood tree was initially inferred after which the inclusion of 1 to 10 gene flow events (-M) between different populations and species were allowed to improve model fit (Supplementary Methods 1). Such migration events represent either population admixture or shared ancestral polymorphism retained after population isolation. For each setting the model was ran for 10 iterations while sites were pooled into blocks of 500 SNPs (-k 500) to account for linkage disequilibrium. The tree was rooted using *C. sayi* and only SNPs without missing population allele frequencies were included (n=12,321,150) to minimize bias in variance-covariance matrix estimation. To delineate the optimal number of migration events we compared the mean log likelihood of a model with that of a model containing one additional migration event using a t-test. Initiating at M=0 we selected the M value for which no significant increase in likelihood could be detected. Tree topology and admixture events were visualized using the internal TreeMix plotting function. In addition, interspecific admixture events identified by TreeMix were formally tested using the f 4statistic ^28^ implemented in the *fourpop* module of TreeMix v1.13 ^27^. Standard errors for f 4 statistics were calculated in blocks of 500 SNPs.

### Structural variation analysis

Patterns of genomic divergence (*F*ST) and outlier analysis revealed that sites with elevated divergence between high- and lowland species were generally clustered into contiguous genomic regions that potentially comprise structural variations (SV) like inversions or translocations. We searched for the presence of SV in all scaffolds containing at least ten consecutive 20kb windows (200kb in total) with an *F*ST > 0.1 in the overall comparison between the resequenced high- and lowland individuals (Fig. 4a). This selection procedure resulted in twelve scaffolds with continuous regions of elevated divergence. These twelve scaffolds contained at least two RADtags with an outlier SNP in one of the within-island high-lowland comparisons (see “Outlier loci detection”), which additionally supports that these regions are associated with high-lowland differentiation in at least one of the islands.

We used BreakDancer v1.3.6 ^72^ to screen for anomalies in the insert size or orientation of read pairs that flanked each genomic region of elevated divergence. More precisely, BreakDancer v1.3.6 was run on all individual bam files and we searched for SV whose breakpoints (i) are located within ±20kb of the boundaries of the region with elevated divergence; (ii) have a maximal quality score of Q = 99 and (iii) are supported by a significantly different sequencing coverage between high- and lowland individuals (Welch *t*-test; *P* < 0.05). Regions matching these criteria were considered as contiguous SV for further analysis.

We tested if the sequence composition at SV correspond to the presence of distinct high- and lowland alleles by means of a PCoA analysis on SNPs located within each SV. If haplotypes at the SV represent distinct alleles, we expect individuals in the PCoA to be clustered into three distinct groups corresponding to individuals homozygote for the highland allele, individuals homozygote for the lowland allele and a group of heterozygote individuals that are situated intermediate between both homozygote groups. To further confirm that the three clusters correspond to individuals with different genotypes for distinct highland or lowland alleles, we calculated average nucleotide diversity (π) at the SV and tested if π is significantly higher in the individuals in the heterozygote cluster compared to those in the two homozygote clusters. PCoA analysis was performed with the adegenet 2.1.3 ^63^ package in R v.4.0.3 using SNPs located within each SV and filtered with VCFtools v0.1.16 ^61^ for a genotype quality > 30 and presence in all 32 resequenced individuals and kept one out of 1000 SNPs to reduce computational time.

We tested if the SV reduced recombination between high- and lowland associated alleles by calculating pairwise *r*^2^ values between SNP genotypes across the entire length of the selected scaffolds and plotted the distribution of SNPs that are in perfect linkage disequilibrium (*r*^2^=1). If the SV suppresses recombination between both alleles, *r*^2^ = 1 values are expected across the entire length of the SV. We further compared patterns of *r*^2^ between this set that includes all individuals and a set that only includes individuals that are homozygous for the allele associated with the lowland ecotype and, thus, expected to show patterns of free recombination. *r*^2^ calculations were performed with VCFtools v0.1.16 ^61^ based on the same vcf file as used for the PCoA analysis (min GQ > 30, genotypes present in all individuals), but additionally filtered for a minimum allele frequency of 0.05 and an additional SNP thinning of either 1/1000 or 1/10,000 to reduce the number of SNPs to less than 500. Suppression of recombination along the SV of interest was further tested by comparing profiles of nucleotide diversity (π) between individuals that are homo-en heterozygous for the SV, wherein heterozygous individuals are expected to consistently show higher nucleotide diversity compared to homozygotes across the entire length of the SV.

Phylogenetic relationships between the haplotypes located at the SV were estimated by maximum likelihood with IQ-TREE ^65^, specifying 1000 ultra-fast bootstrap samplings ^66^. Before tree estimation, best fitting substitution models for each window were selected using ModelFinder ^67^ as implemented in IQ-TREE. We only included individuals that are homozygous for the SV to estimate phylogenetic relationships because phasing errors in individuals that are heterozygote for the highly divergent alleles may lead to erroneous recombinant haplotypes and highly inaccurate phylogenies ^73^.

We compared the timing of the divergence between high- and lowland alleles across the different SV using the divergence between the *Calosoma* species from the Galapagos and the mainland species *C. sayi* as relative calibration point. Analyses were performed with BEAST v2.6.0 ^74^, specifying the best fitting substitution model for each SV as selected by ModelFinder in our IQ-TREE analysis, a random local clock model, empirical base frequencies, and Yule tree prior. We further specified a clade with individuals with the highland allele and a clade with individuals with the lowland allele, which were both constrained to be monophyletic if supported (>95%) by our maximum likelihood analysis. Analyses were run for 50M generations and we only used samples from the stationary phase of the Markov chain, comprising at least 25M generations, for consensus tree construction and divergence time estimation. Mean and 95% HPD of the relative timing of the split between the clades comprising individuals with the high- or lowland alleles was then divided by the relative timing of the split between *C. sayi* and the Galapagos species. To compare the relative divergence time between both alleles among the different SVs. Relative divergence of this split between high-and lowland alleles was compared with the relative divergence time of the different species in this radiation. Timing of the split of these highland species, relative to the divergence from the outgroup *C. sayi*, was estimated by running a multispecies multilocus coalescence analysis with *BEAST ^74^ based on a random selection of 50 windows of 20kb (see Phylogeny in Methods section for details on the extraction of fasta files for these 20kb windows) that were located outside the SV. We only selected those windows for which ModelFinder reported a HKY substitution model as this allowed us to specify the same HKY substitution model for all 50 windows simultaneously in the *BEAST analysis. Like our previous analysis on individual SV, we specified a random local clock model, empirical base frequencies, and a Yule tree prior.

## Acknowledgements

This study is largely based on specimens obtained from the extensive collection of the late Konjev Desender (†), who collected most *Calosoma* individuals used in this study. Wouter Dekoninck, Charlotte De Busschere and Henri W. Herrera provided help in collecting additional specimens. Outgroup specimens were kindly provided by Martin Husemann. We thank Mado Berthet for designing the pictograms of the *Calosoma* species, Alain Drumont for preparing the specimens and Camille Locatelli for help in taking pictures of the specimens. We are indebted to Jenna Mann and Cathlene Eland for demonstrating and sharing the RADseq protocol. Field logistic support was provided by the Charles Darwin Research Station (CDRS; Isla Santa Cruz, Galápagos, Ecuador); the Galápagos National Park Service and the Department of Forestry, Ministry of Agriculture of Ecuador. Financial support for the expeditions was achieved from the Royal Belgian Institute of Natural Sciences and the King Leopold III Fund. Analyses were carried out using the STEVIN Supercomputer Infrastructure at Ghent University, funded by Ghent University, the Flemish Supercomputer Center (VSC), the Hercules Foundation and the Flemish Governement—department EWI. This work was financially supported by the Belgian Science Policy - BelSPo (BRAIN-Be projects BR/121/PI/GENESORT and BR/175/PI/PARAWINGS).

