## Supplementary Information for "Progressive spread of chromosomal inversions blends the role of colonization and evolution in a parallel Galápagos beetle radiation"

Supplementary Tables S1-S5

Supplementary Figures S1 –S3

Supplementary Methods 1: “Admixture analysis”

**Table S1 |** Overview of the specimens used for genome assembly, restriction-site associated DNA (RAD) sequencing and whole genome resequencing.

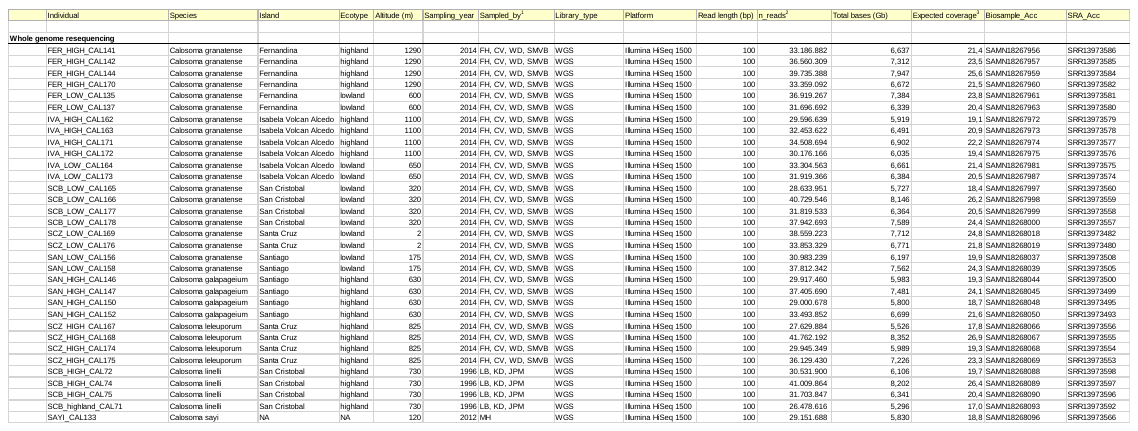

**Table S1 (continued)**

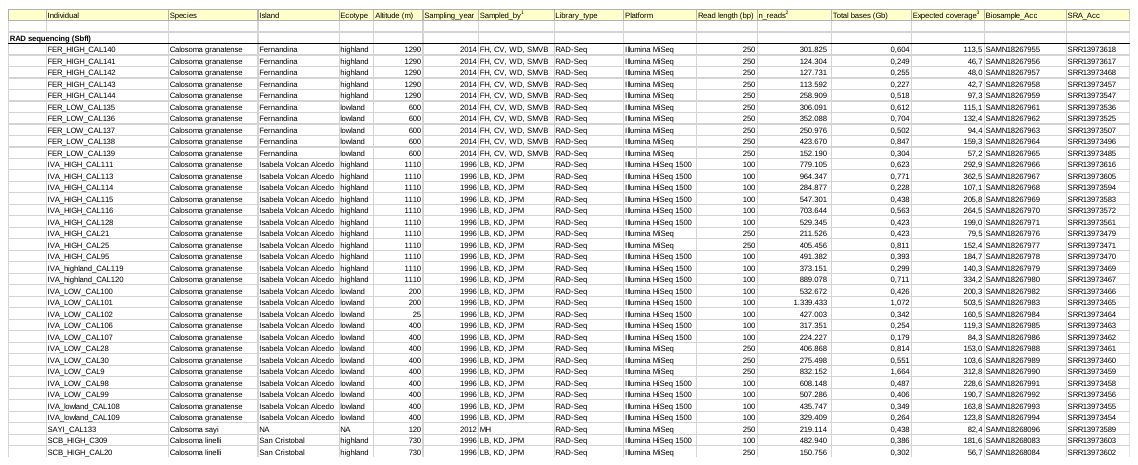

**Table S1 (continued)**

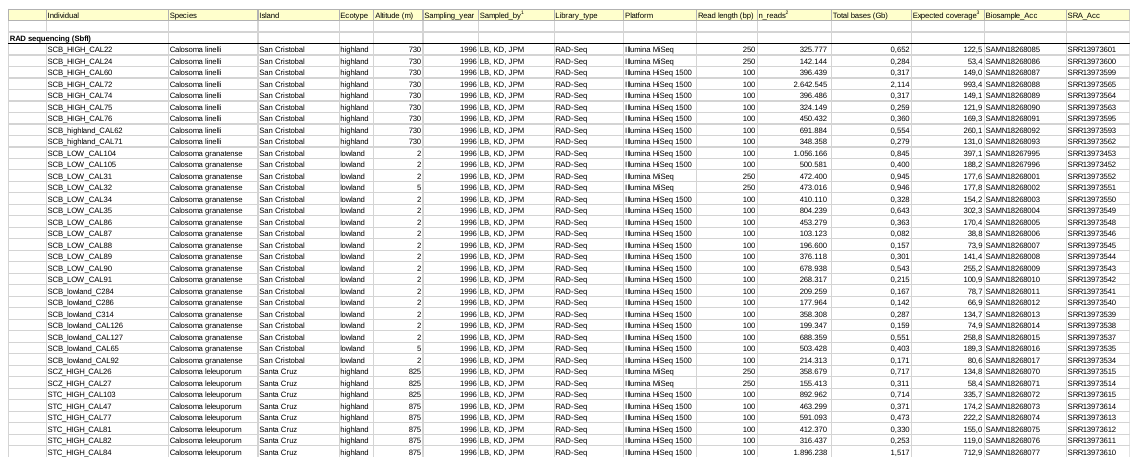

**Table S1 (continued)**

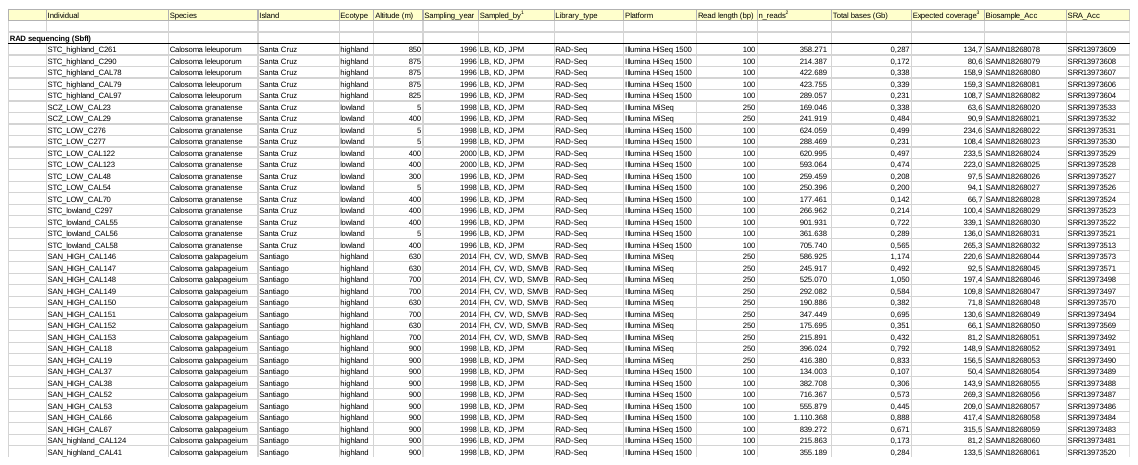

**Table S1 (continued)**

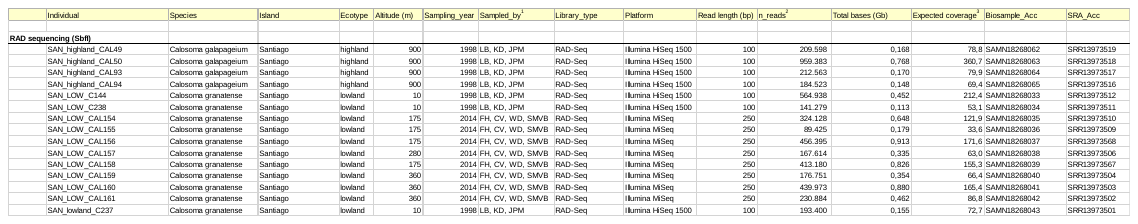

**Table S1 (continued)**

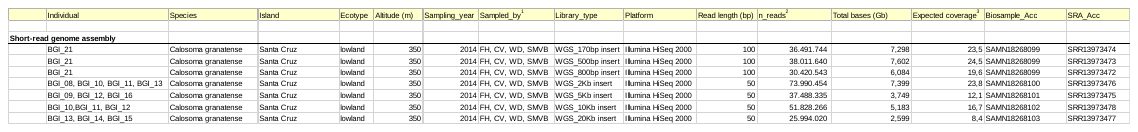

**Table S1 (continued)**

Notes:

^1^ LB=Léon Baert,WD = Wouter Dekoninck,KD = Konjev Desender, FH = Frederik Hendrickx, MH = Martin Husemann, JPM = Jean-Pierre Maelfait, CV = Carl Vangestel, SMVB= Steven M Van Belleghem

^2^ Paired reads

^3^ Coverage of RADseq data represents expected coverage of forward reads and 1,330 SbfI restriction sites in the genome assembly

^4^ Coverage of WGS data is based on an expected genome size of 0.310Gb (k-mer based estimate with k-mer size of 15)

**Table S2 |** Summary of the number of individuals of each population or species that were used for RAD sequencing and whole genome resequencing.

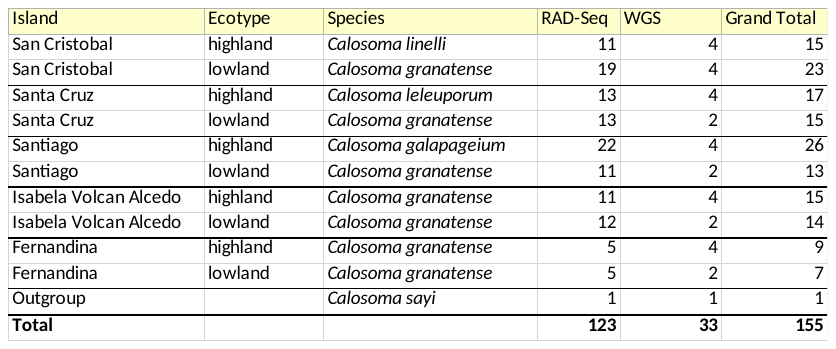

**Table S3 |** Assembly statistics and repeat content of the *Calosoma granatense* genome assembly

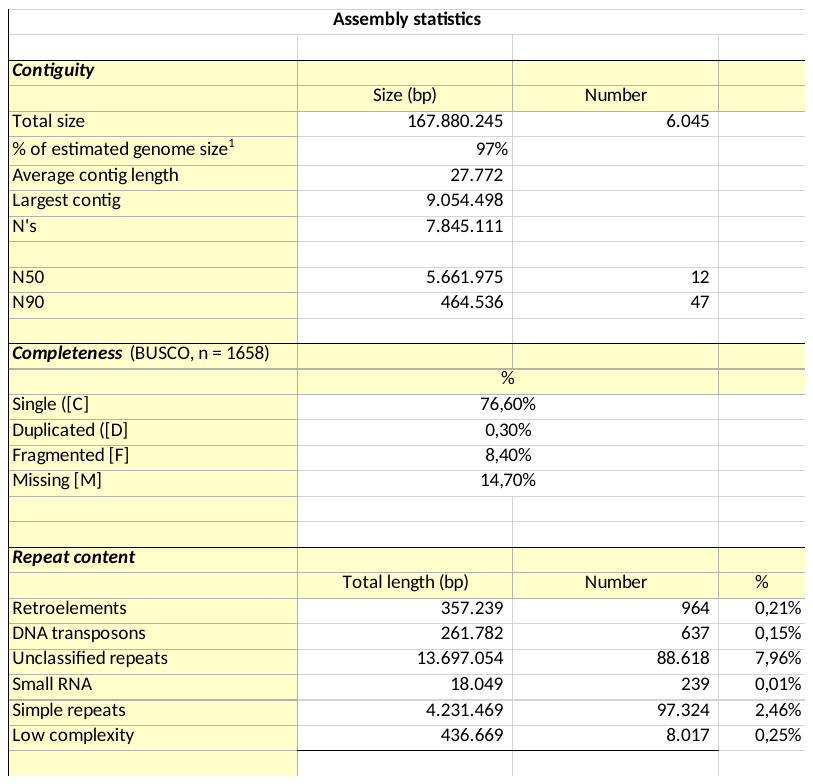

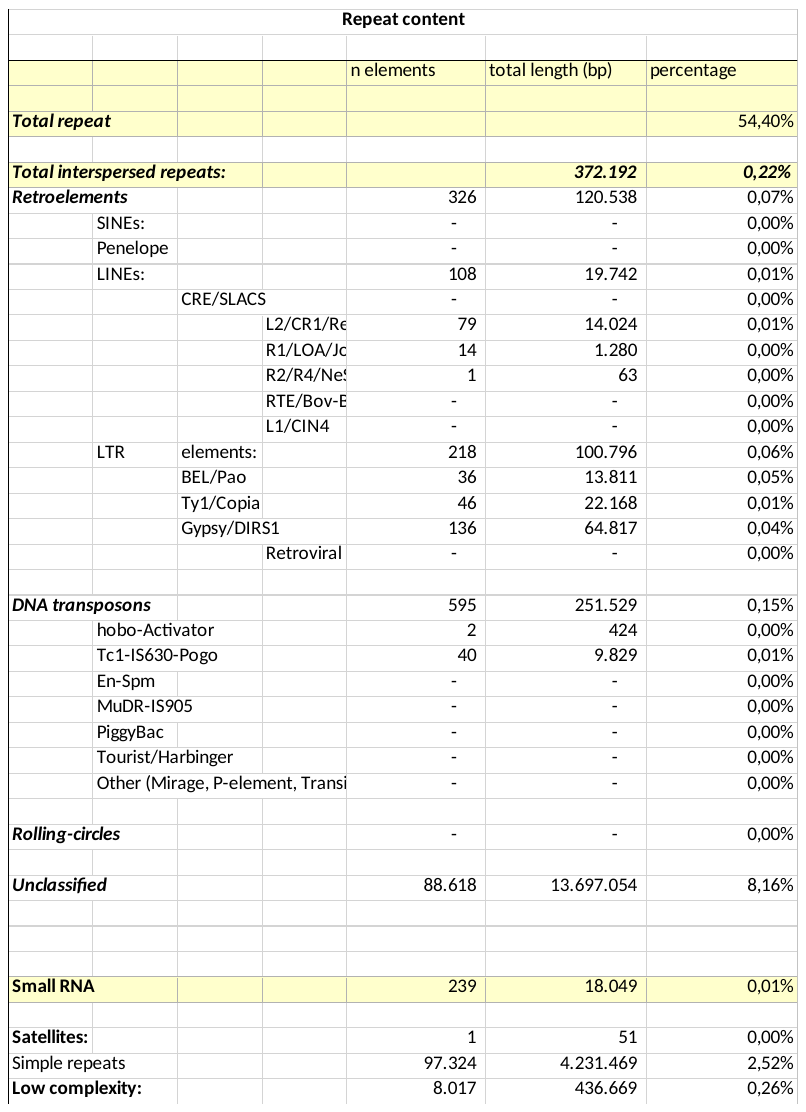

**Table S4 |** Detailed representation of admixture events identified by *TreeMix* v1.13. Type: intra- or interspecific admixture. Weight and migration edge: weights of inferred gene flow events (migration edge) in modified Newick format. The following columns depict results from the f_4_ analysis. f_4_ test: the four population tree. f_4_ value, SE and Z: the f_4_ estimate, its standard error and associated Z score (f_4_ estimates with an associated absolute value of a z-score larger than 2.58 were regarded as significant deviations from their null value (no admixture) and are indicated in bold). Last column provides a brief description of interspecific admixture event that is being tested.

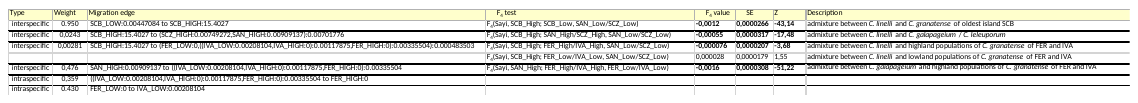

**Table S5 |** Summary of the SV detected by *BreakDancer* v1.3.6 that overlap with the largest genomic regions with elevated divergence. Scaffold: scaffold with the inversion. SV type, start, end and length: type, start and end of the SV reported by *BreakDancer*. Q: quality-score of the SV reported by *BreakDancer*, with Q = 99 being the maximal score. *P*_high-low_: *P*-value (Welch-test) of the difference in the number reads that support the inversion between high- and lowland individuals. *n* RADtags: number of RADtags with an outlier SNP that is shared in at least two within-island ecotype comparisons. *P*_geno_, *P*_HH-HL_, *P*_HH-LL_ and *P*_HL-LL_: *P*-value (one-way ANOVA) of the difference in nucleotide diversity at the SV between individuals identified as homozygous highland (HH), homozygous lowland (LL) and heterozygous (HL) and the respective pairwise comparisons between these genotypes by mean of a Tukey HSD test. *r*_S_(HH) and *r*_S_(LL): Spearman rank order correlation between island age and nucleotide diversity of individuals homozygous for the highland allele (HH) or lowland (LL) allele. Next columns show corresponding *P* -values of the correlation.

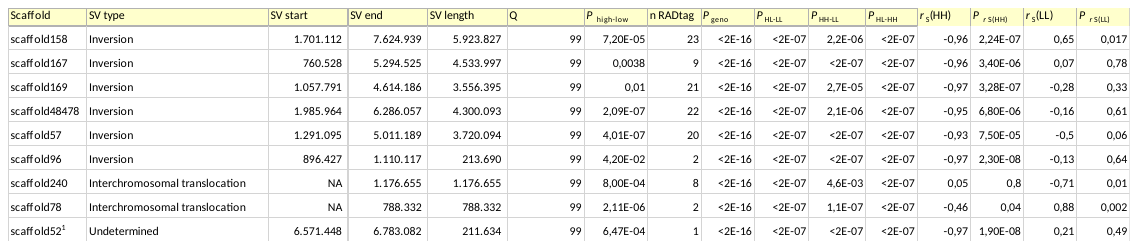

Notes

The region of elevated divergence on this scaffold was flanked by a deletion at position 6,571,448 and a deletion at position 6,783,082 that were both statistically supported by a significantly different number of reads between high- and lowland individuals (P high-low = 0.000647 and P high-low = 0.0000125 respectively).

**Fig. S1.** – **Structural variations (SV) underlie distinct alleles associated with high-lowland divergence.** Each column shows the result of a single scaffold on which the SV was located. **First row**: *F*_ST_ distribution (20kb windows) based on a comparison between all high- versus lowland individuals. Blue bottom line shows the location of the chromosomal inversion detected by *BreakDancer* ^72^. Red dots show the location of RADtags identified as outlier loci in at least one within-island highland-lowland comparison. **Second row:** Location of SNPs in perfect linkage disequilibrium (*r*² = 1). Grey dots above diagonal show *r*² = 1 values for all 32 resequenced individuals. Grey dots below diagonal show *r*² = 1 values for homozygous LL individuals only. **Third row:** PCoA based on SNPs located at the inversion (blue line in upper row panels). HH, LL and HL refer to the cluster of individuals genotyped at the inversion as homozygous for the highland allele (HH), lowland allele (LL) and heterozygous (HL). **Forth row**: Differences in nucleotide diversity at the inversion between individuals genotyped as HH, HL and LL in the third row panels. **Fifth row**: Maximum likelihood tree of the nucleotide sequence at the inversion excluding individuals genotyped as heterozygotes (HL). Node values represent bootstrap values based on 1000 replicates. The tree was rooted with the mainland species *C. sayi*. **Sixth row**: Relationship between individual nucleotide diversity at the inversion and the progression of the islands. Only individuals genotyped as homozygous for the lowland allele (LL, yellow) and highland allele (HH, remaining colors) are included. Color codes are as in Fig. 2.

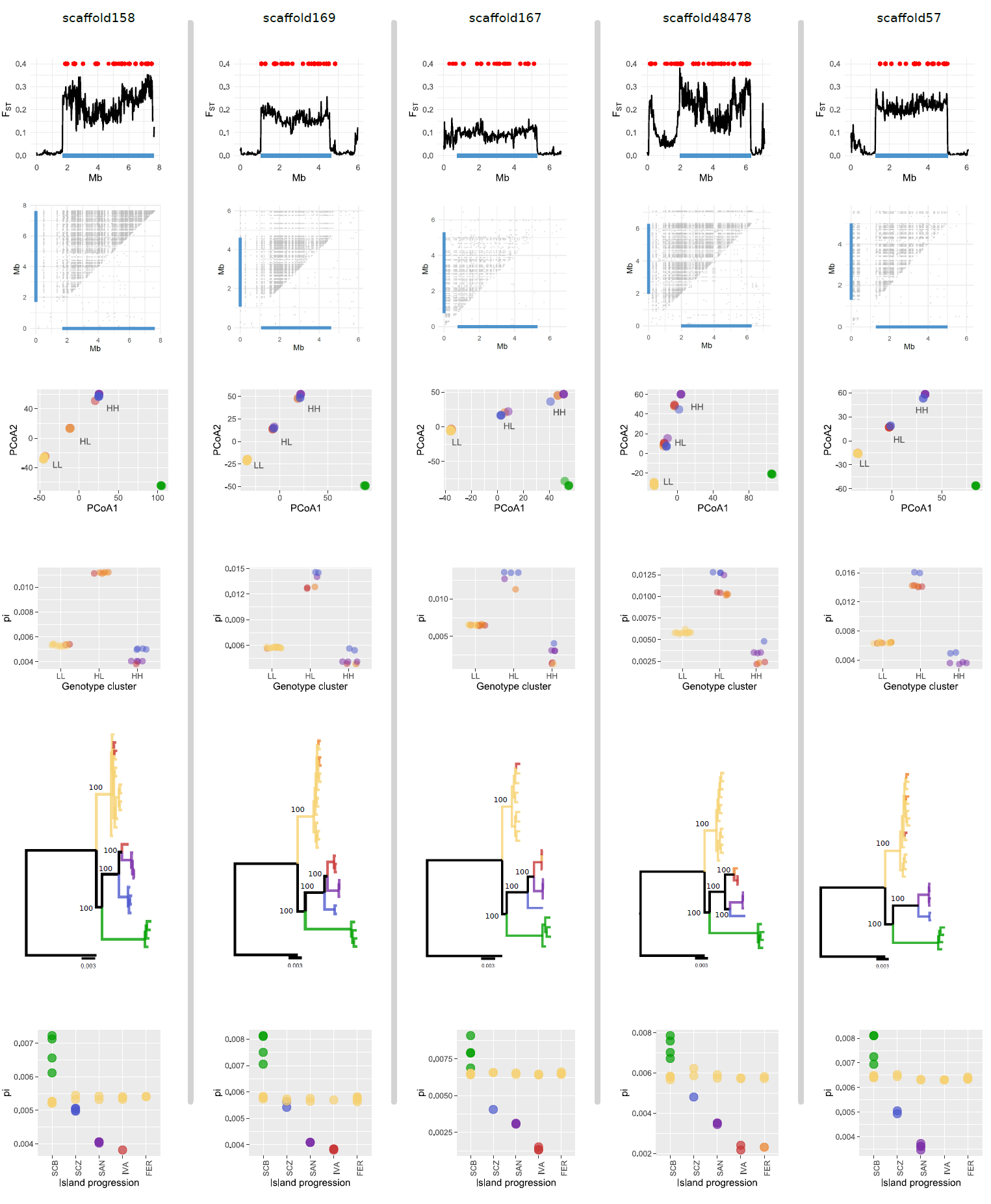

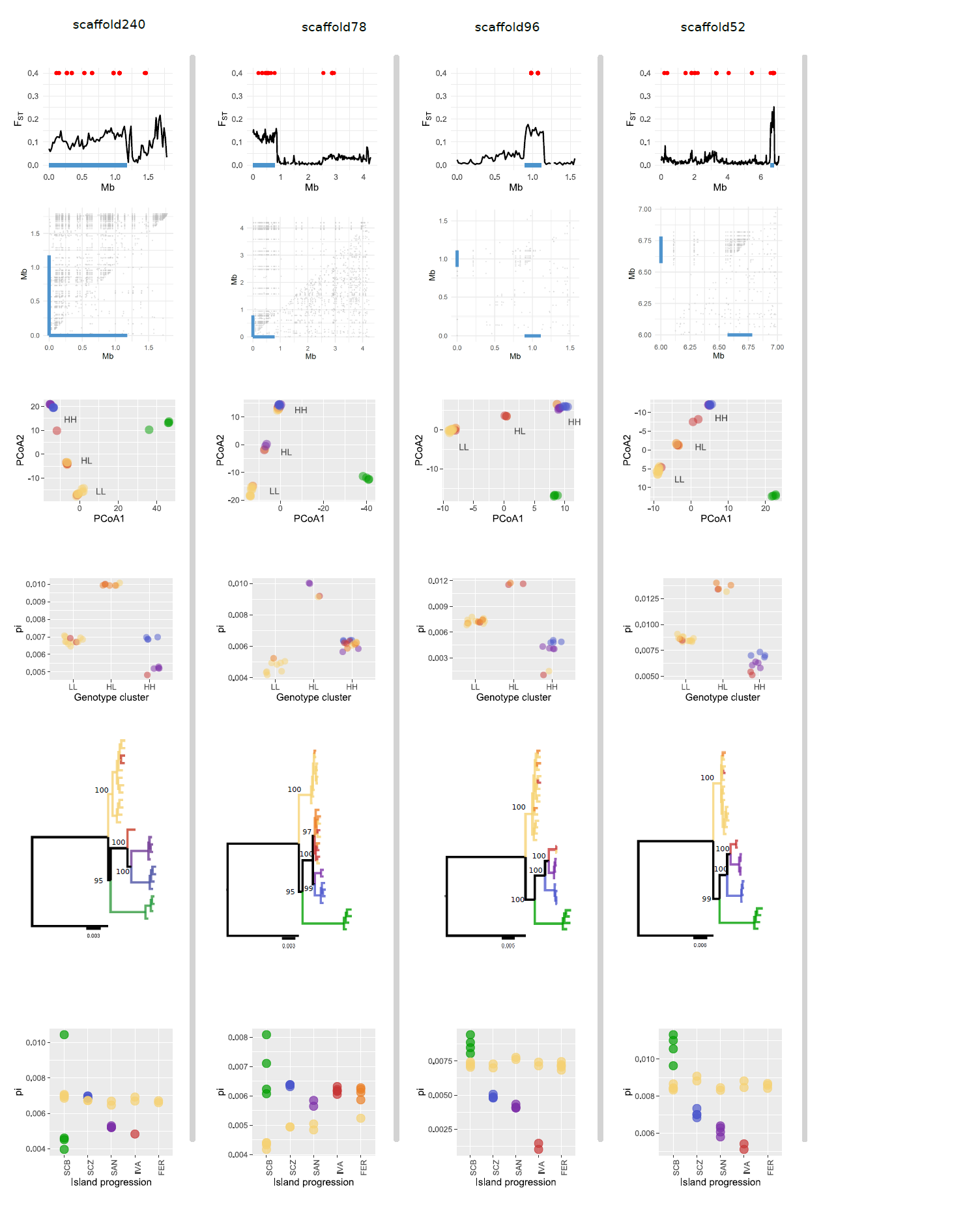

**Fig. S2.** – **Patterns of genetic differentiation and nucleotide diversity at scaffolds with structural variations associated with highland-lowland divergence.**

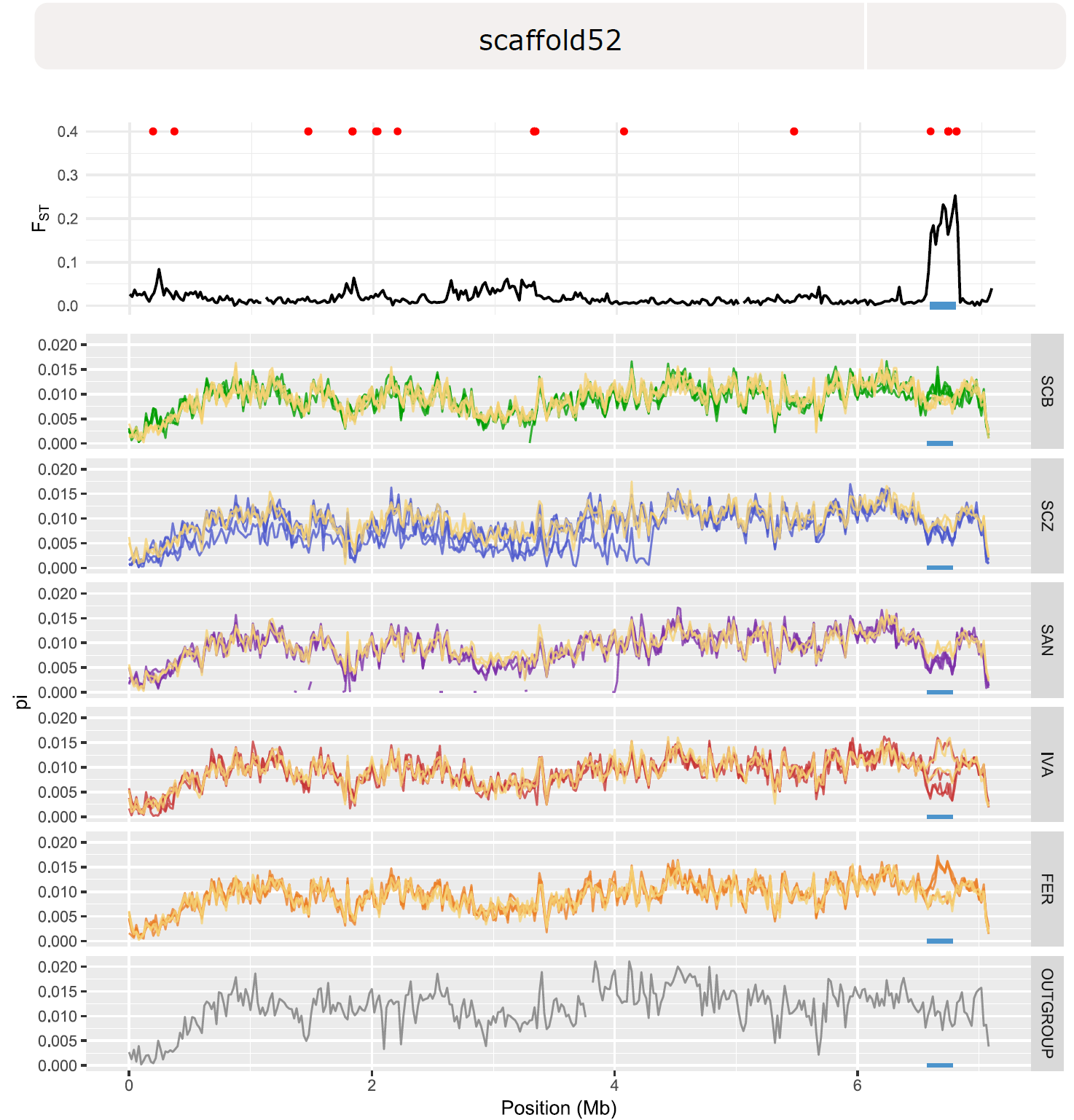

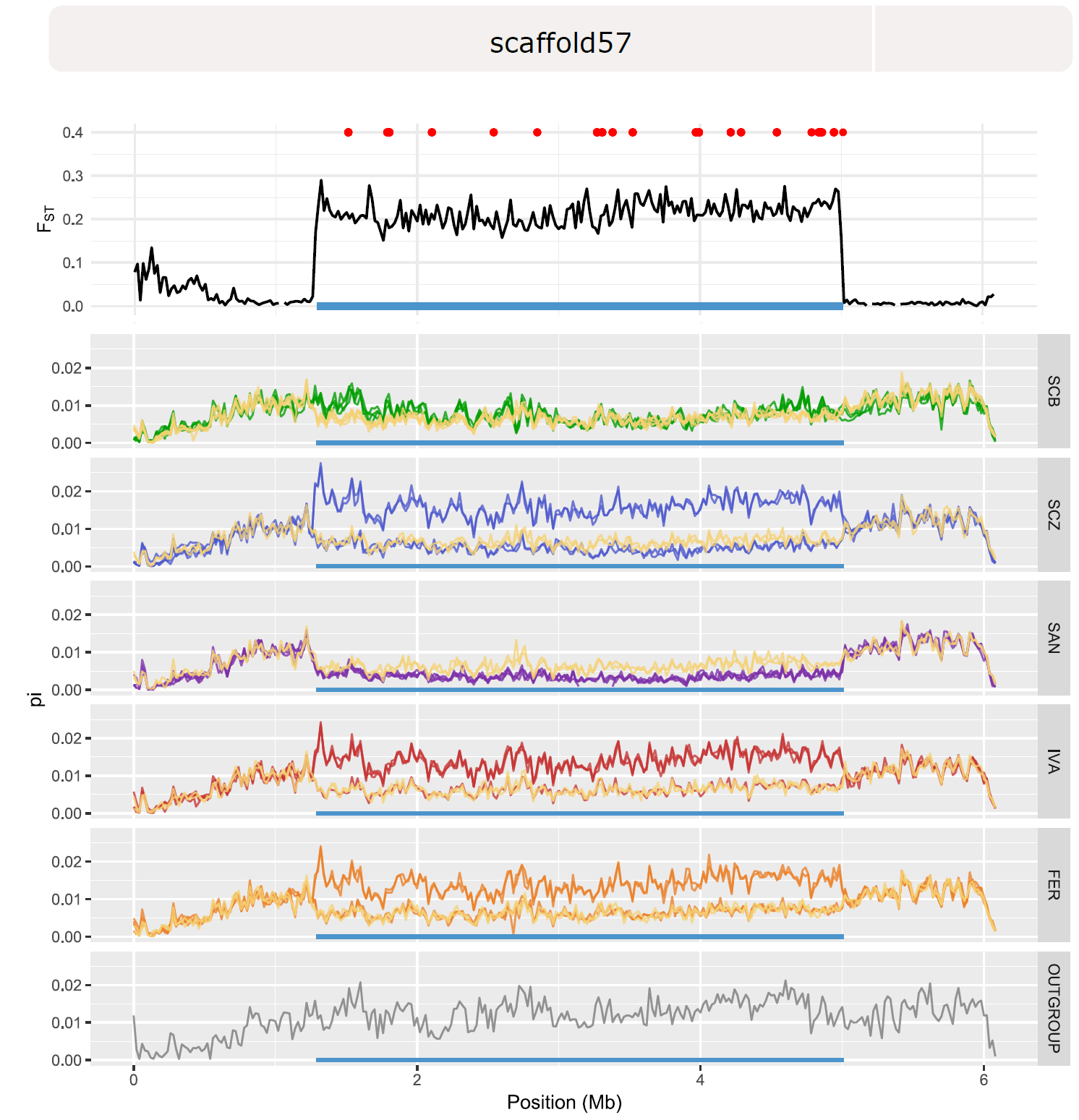

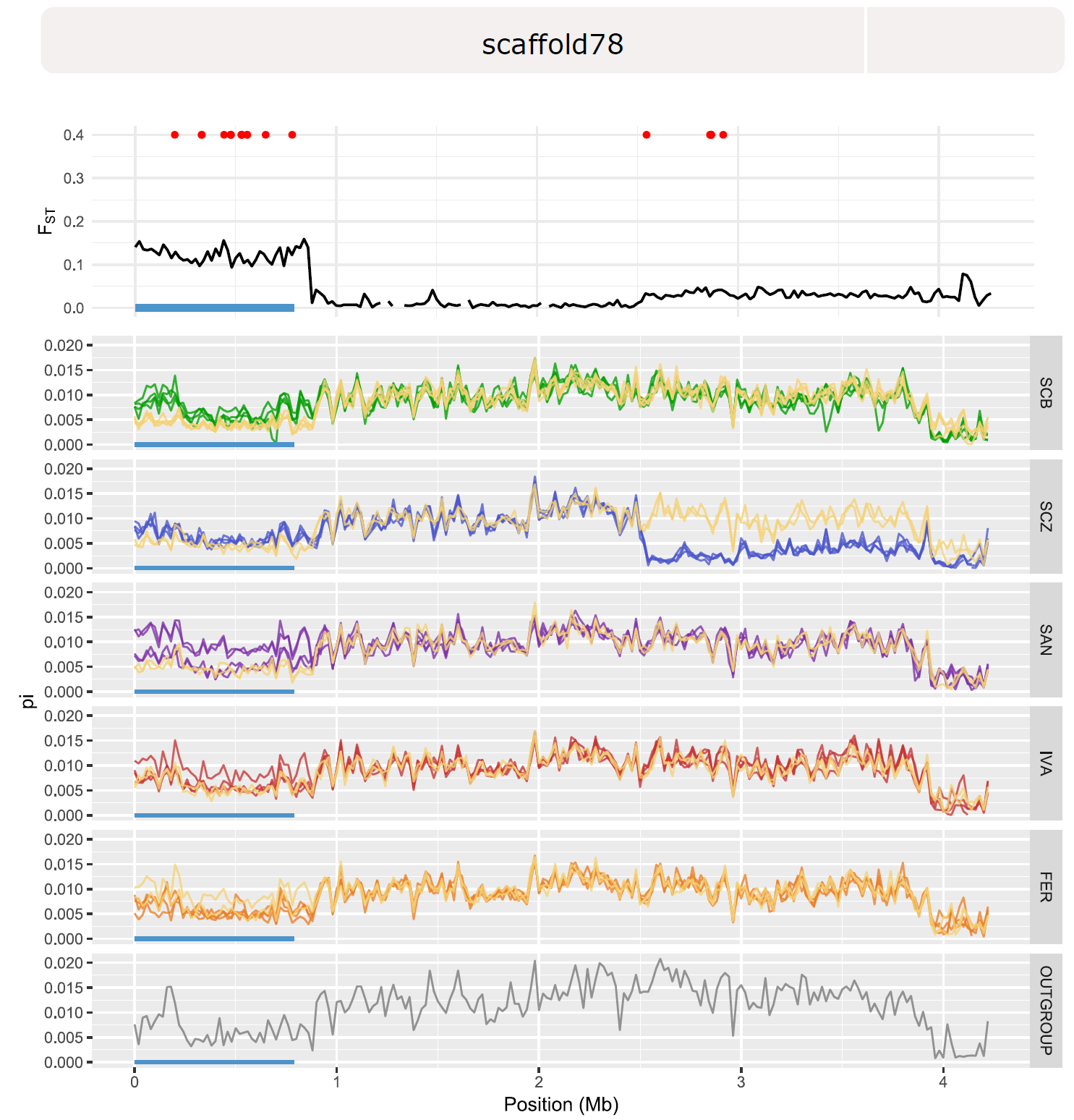

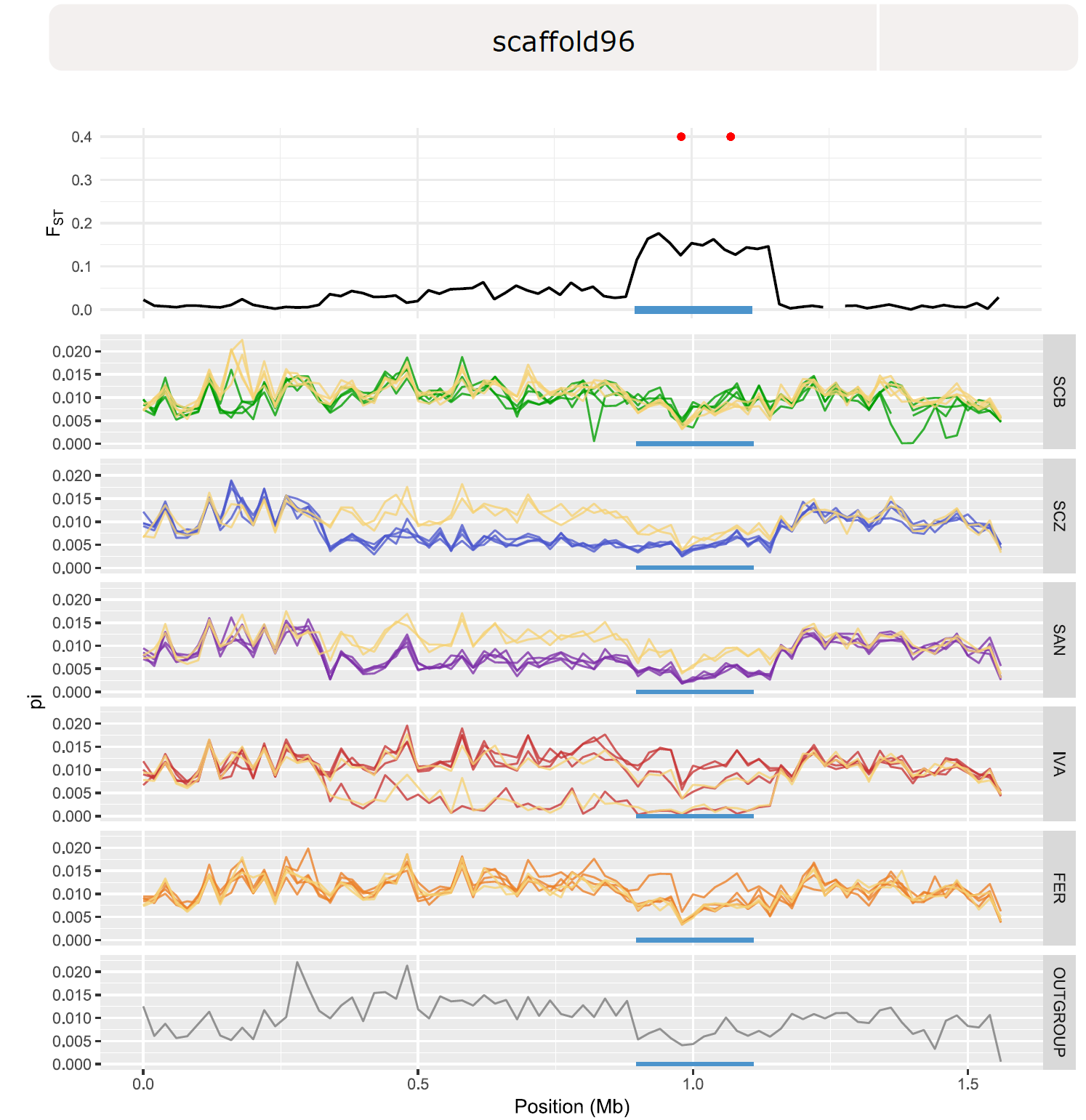

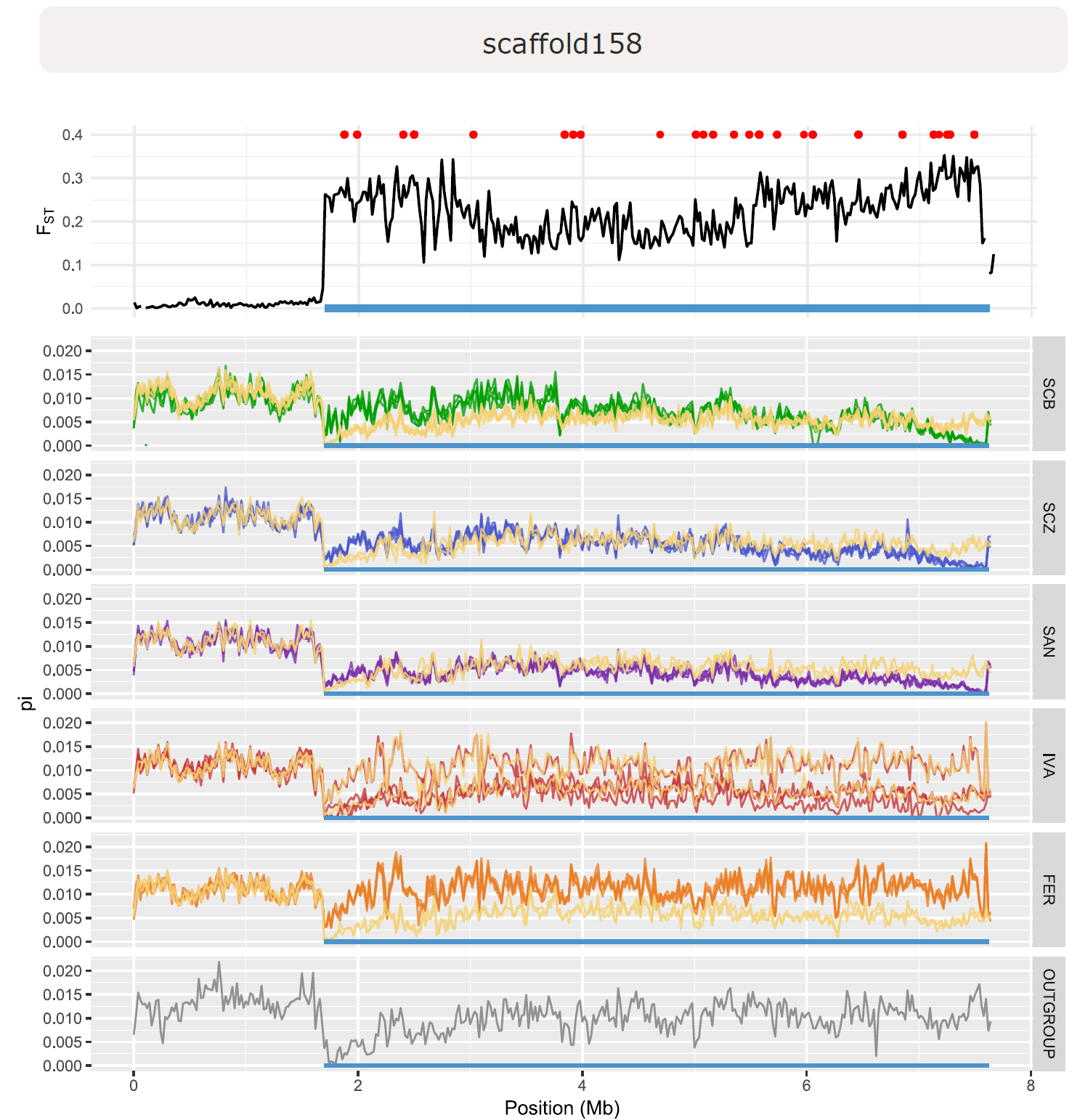

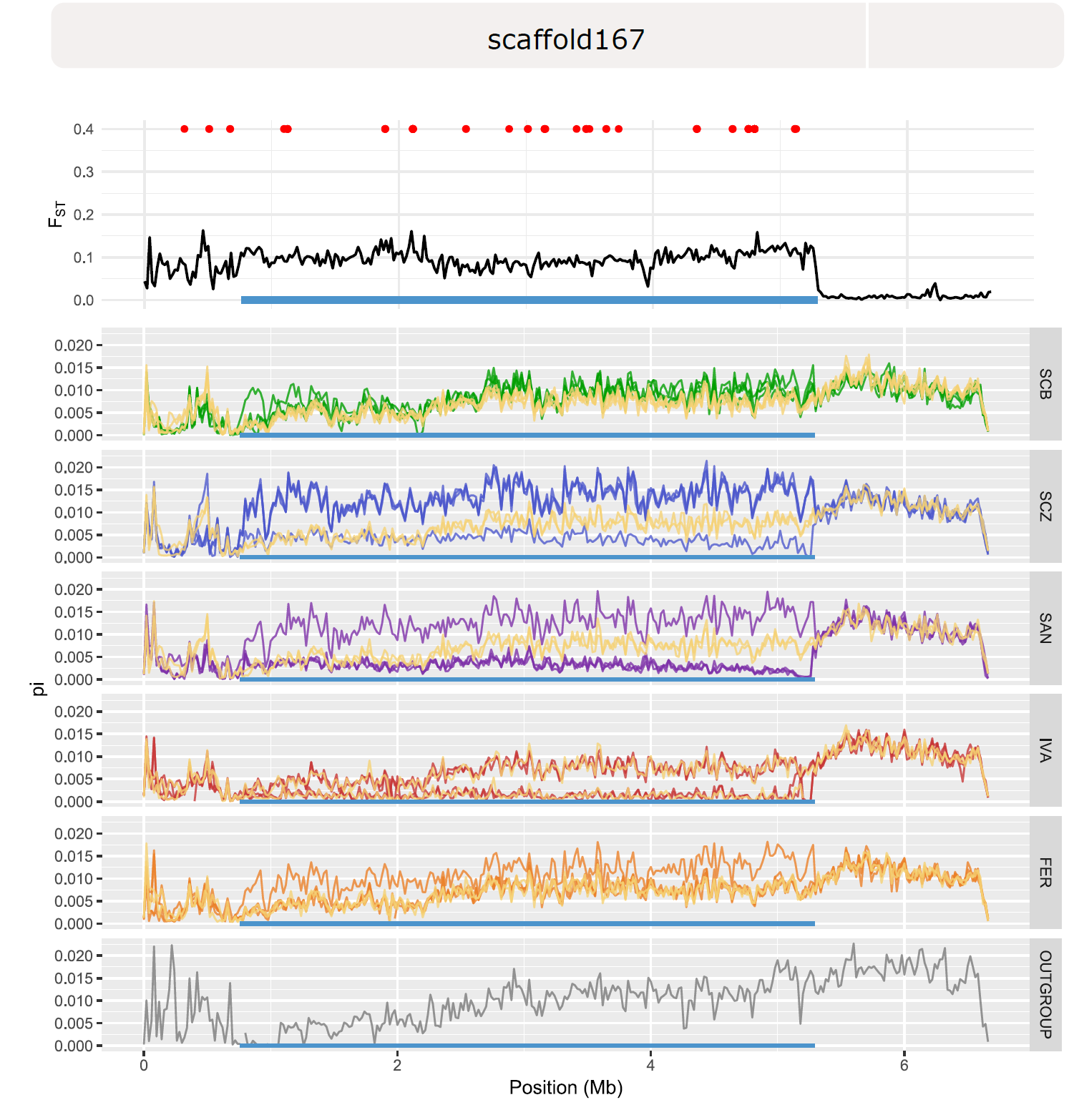

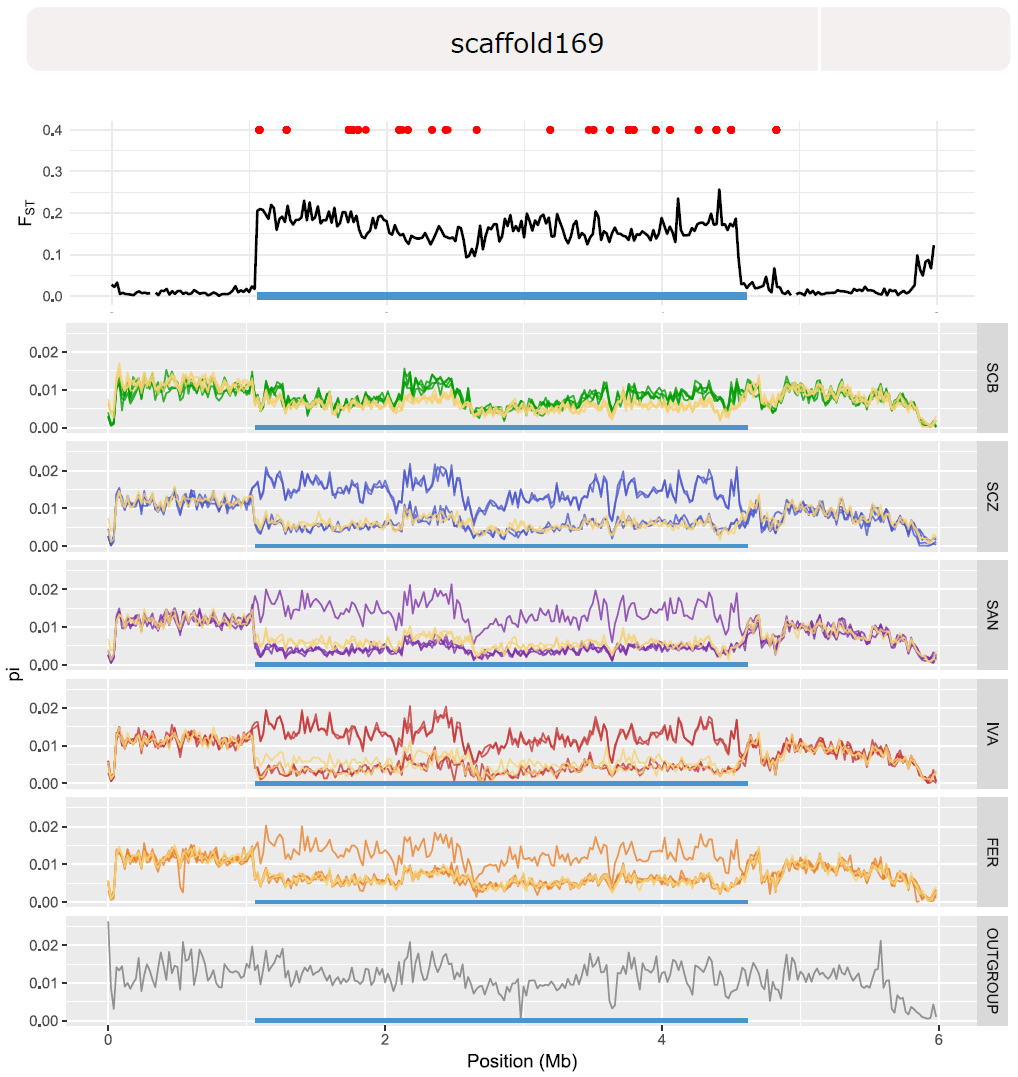

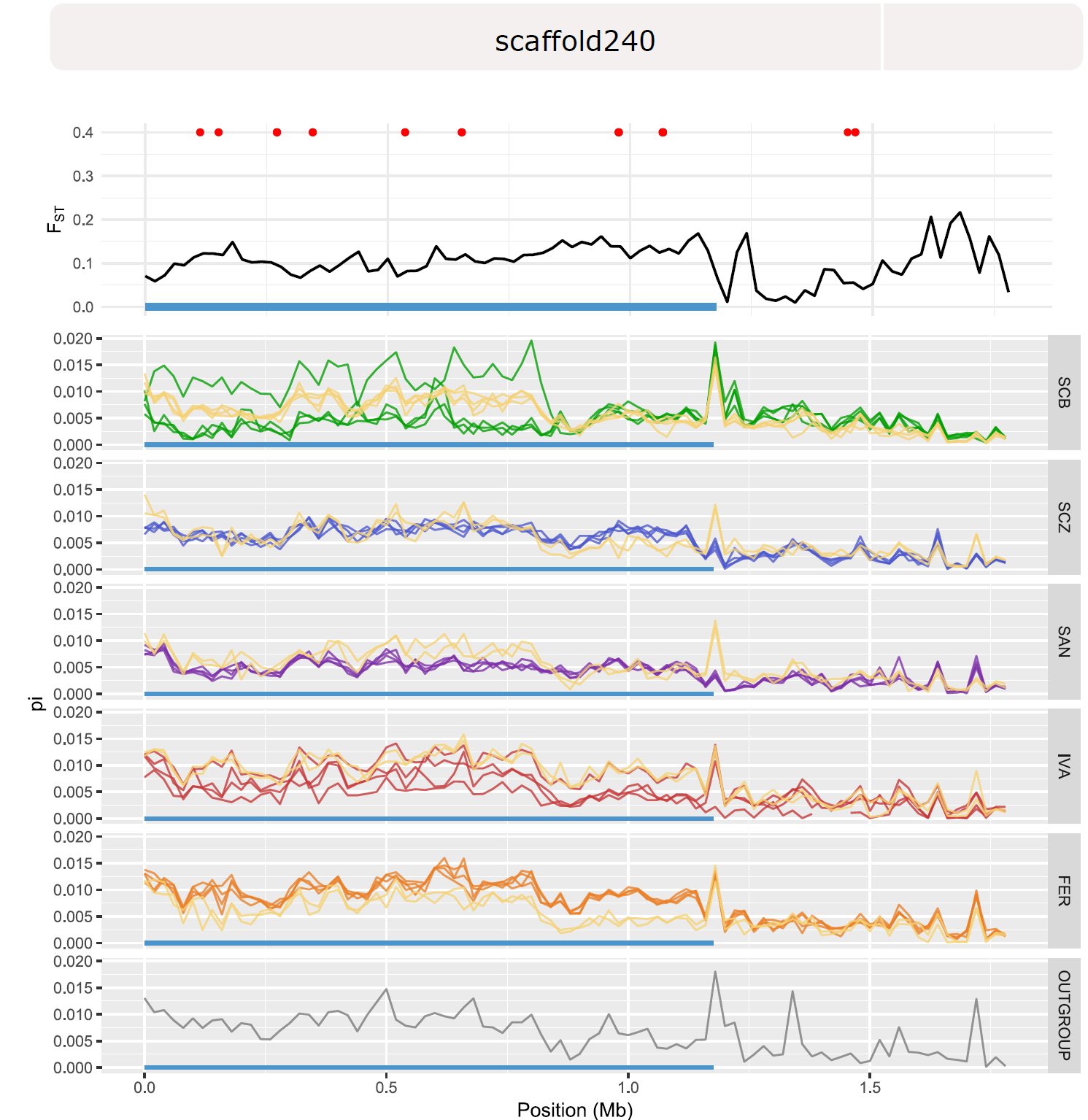

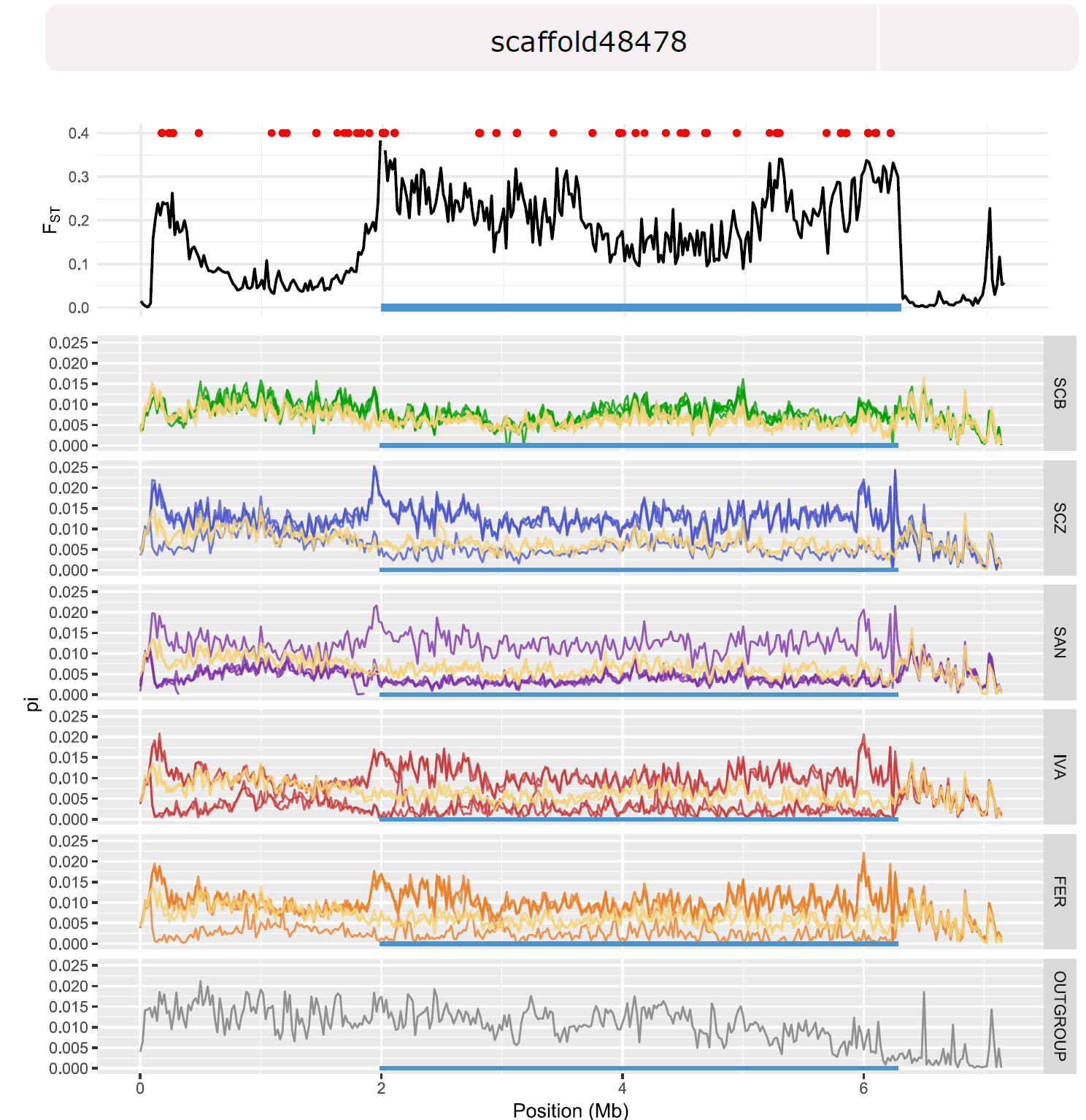

**Fig. S3.** – **Segregation patterns of high- and lowland alleles at the nine investigated SV associated with high-lowland divergence.** Yellow squares represent individuals homozygote for the lowland allele; grey squares are individuals heterozgote for the high and lowland allele; remaining colored squares are individuals homozygote for the highland allele, with the colors corresponding to the highland population and species colors shown in Fig. 2.

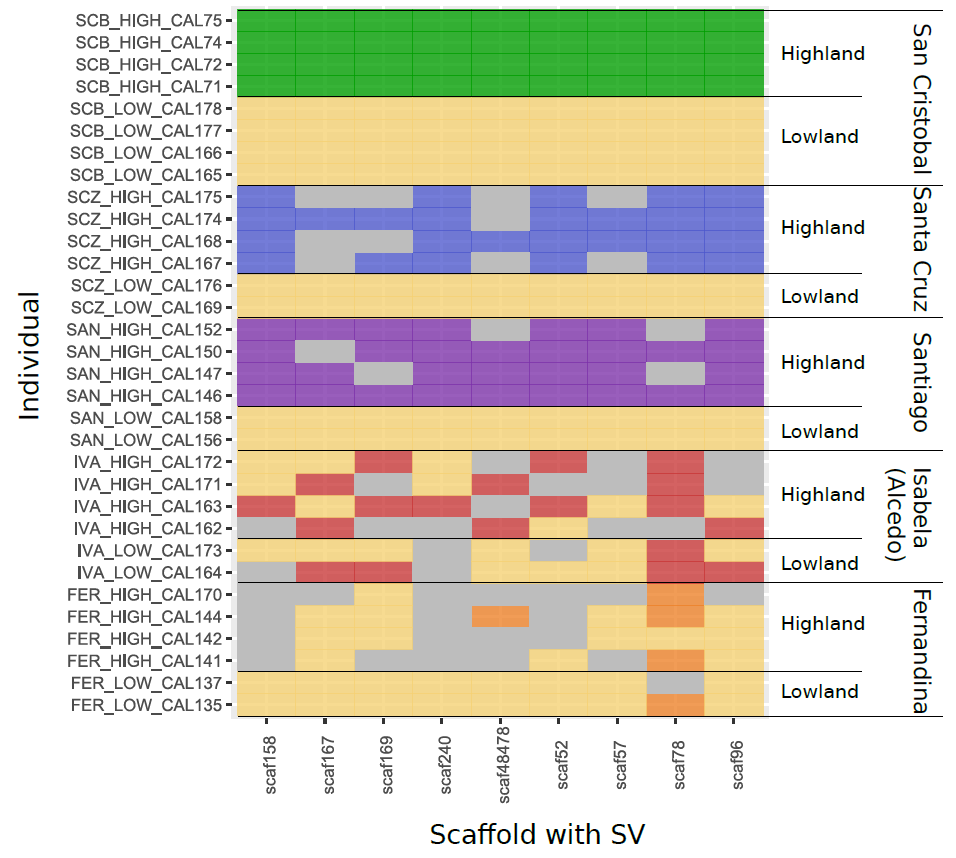

**Supplementary methods 1: “Admixture analysis”**

We used TreeMix v1.13 (1) to estimate genetic relationships and admixture between 32 resequenced Calosoma specimens sampled across the archipelago and included *C. Sayi* as an outgroup. After inferring an initial maximum likelihood tree we allowed up to 10 migration events to improve the data fit. For each setting we ran the model for 10 iterations. As adding complexity to the model (i.e. migration events) will increase the likelihood, we attempted to delineate the optimal number of migration events as the one for which the mean likelihood reached a plateau, i.e. when the mean likelihood did not significantly increase when comparing it with that of a model containing 1 additional migration event. Pairwise t-tests indicated that at 6 migration events such a plateau was reached as the mean log likelihood of a model with 7 migration events did not significantly differ from one with 6 migration events (t_18_=1.497, p=0.15) (Fig. 1).

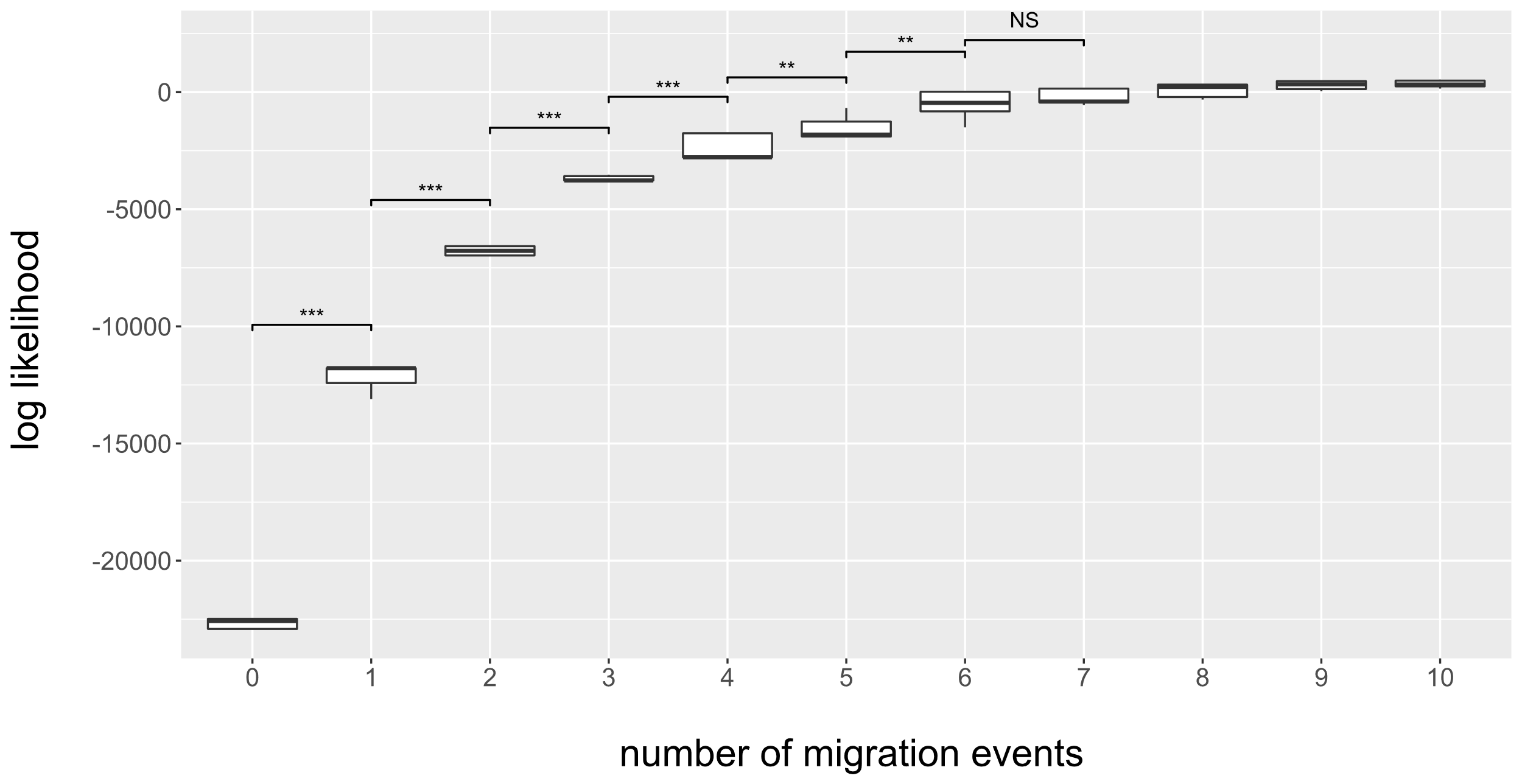
 **Figure 1 | Likelihood profile of TreeMix models with increasing number of migration events.** Plot depicts an increasing median log likelihood as more migration events are added to the model. Symbol above bracket connecting two adjacent number of migration events indicates the p-value of a t-test between their mean log likelihood (* p<0.05, ** p<0.01, *** p<0.001, NS p>0.05).

The corresponding tree suggested interspecific admixture or retention of ancestral variation between the highland species *C. linelli* (SCB High) inhabiting the oldest island and respectively i) the other highland species *C. galapageium* (SAN High) and *C. leleuporum* (SCZ High), ii) the lowland species *C. granatense* residing on the oldest island (SCB Low) and iii) an ancestor of *C. granatense* that gave rise to the lineages found on the youngest islands Isabela and Fernandina. Additional evidence for interspecific admixture was found between the highland species *C. galapageium* (SAN High) and the high- and lowland populations of *C. granatense* inhabiting the youngest islands (FER High, IVA High, IVA Low). Finally, TreeMix results indicated intraspecific admixture between the *C. granatense* populations residing on the youngest islands Fernandina and Isabela (Table S4).

We further explored admixture events across the archipelago by formally testing interspecific migration using F_4_ statistics (2,3). When no admixture takes place, the F_4_(*a*,*b*;*c*,*d*) statistic should result in differences in allele frequencies between population *a* and *b* that are uncorrelated to those observed between population *c* and *d*, i.e. F_4_(*a*,*b*;*c*,*d*)=0. We assigned the outgroup *C. sayi* to population *a*, assuming that no admixture has taken place between this outgroup and any of the Calosoma species or populations of the Galápagos archipelago, such that a significant positive f_4_ statistic would indicate admixture between populations *b* and *d*, and a significant negative f_4_ statistic would point towards admixture between populations *b* and *c*. When testing the interspecific admixture events previously identified by TreeMix, we pooled samples according to the tree topology and migration events, i.e. specimens belonging to respectively i) *C. leleuporum* and *C. galapageium*, ii) the highland populations of Isabela and Fernandina, iii) the lowland populations of Isabela and Fernandina, and iv) the lowland populations Santa Cruz and Santiago. F_4_ statistics were estimated using the *fourpop* module of TreeMix v1.13 (1) and associated standard errors were calculated in blocks of 500 SNPs.

Results from the f_4_ analysis corroborated those indicated by the TreeMix graph (Table S4). Ancient variation of *C. linelli* (SCB High) could be traced back into the genomes of the other two highland species *C. leleuporum* and *C. galapageium*, the lowland species *C. granatense* of the oldest island and the highland populations of *C. granatense* of the youngest islands. The analysis further highlighted the absence of such admixture signatures between *C. linelli* and the lowland populations of *C. granatense* found on the youngest islands. In congruence with TreeMix analysis, we found statistical support for admixture between *C. galapageium* and the highland populations of *C. granatense* of the youngest islands.
